# A force-balance model for centrosome positioning and spindle elongation during Interphase and Anaphase B

**DOI:** 10.1101/2021.10.04.463000

**Authors:** Arittri Mallick, Apurba Sarkar, Raja Paul

## Abstract

A computational model in one dimension is proposed to position a single centrosome using astral microtubules (MTs) interacting with the cell cortex. The mechanism exploits mutually antagonistic pulling and pushing forces arising from the astral MTs’ binding to cortical dynein motors in the actin-rich cell cortex and their buckling while growing against the cell cortex, respectively. The underlying mechanism is extended to account for the elongation and positioning of the bipolar spindle during mitotic anaphase B. Besides astral MTs, the model for bipolar spindle involves interpolar microtubules (IPMTs). The composite model can predict spindle elongation and position under various circumstances. The outcome reveals that the bipolar spindle elongation, weakened by decreasing overlap between the antiparallel IPMTs in the spindle mid-zone, is recovered by the astral MTs. The one-dimensional models are extended in two dimensions to include the effect of cortical sliding of the astral MTs for studying the dynamics of the interphase centrosome and the anaphase B spindles in elongated cells. The results reveal that the dynamics in two dimensions stay qualitatively similar to the one dimension.

## I. INTRODUCTION

Microtubules (MTs) form the basis of cellular architecture and dynamics that give rise to complex phenomena with significant implications in cell biology. The MTs can emanate from various cellular machinery and are very useful mechanical propellers of various cellular processes [1–7]. They interact with associated cellular components like molecular motors: both uni-polar and bipolar; the cell cortex, and even other MTs [8–10]. MTs are known to be actively involved in mechanical interactions that are crucial for critical cellular processes like centrosome positioning at the cell center during interphase [2, 11–17], bipolar spindle elongation and positioning during mitotic anaphase B [18–20] etc. Given that MTs play the chief mediator in these processes, it is essential to develop a computational model for studying these processes in great detail.

The centrosome is actively positioned at the cell center during the Interphase of the cell cycle [2, 11, 12, 16]. It is subjected to a force field of pulling and pushing forces that are mediated by cortical motors and centrosome nucleated microtubules, respectively [2, 11, 12]. The centrosome, which is thought to be the primary microtubule-organizing center(MTOC) in mammalian cells, can give rise to microtubules that grow isotropically toward the cell cortex [8]. These microtubules are called astral microtubules. Their dynamic instability parameters ensure the growth till the cell cortex [21] where their interaction with the cortical wall acts as the source of various force generating mechanisms. The astral MTs reaching the cell cortex can associate with the cortical dynein motors that are already bound to the actin-rich cortical layer. Association of the astral MTs with the dynein motor is favored as these motors are minus-end directed. Minus-end directed motion of the motors generate pulling force (toward the cell cortex) on the centrosome through the astral MTs [11]. The event of buckling of MTs against a rigid barrier, when they are grown from a movable source, has long been established [22] and is known to generate a net push on the source, directed away from the rigid barrier. The astral MTs have been found to buckle against cell organelles and cortex to generate pushing force on the centrosome [11]. Eventually, a balance between these forces causes the centrosome to position itself at the cell center. During Interphase, fission yeast *Schizosaccharomyces pombe* (*S. pombe*) employs a similar mechanism to retain the nuclei at the cell center, ensuring that the metaphase plate forms close to it [23]. *S. pombe* is an elongated cell with antiparallel bundles of microtubules arranged along the cell’s long axis [23, 24]. Their minus ends appear to cluster on the nuclear envelope at places near its north and south poles [23]. The plus ends of these microtubules grow parallel to the long axis and bend against the membrane as they try to grow against the cell tips, producing transitory pushing force that eventually positions the nucleus at the cell’s center [23].

The centrosome gets duplicated before the onset of mitosis [25, 26], aiming to build a bipolar array of microtubule-based machinery, called the mitotic spindle, which forms during metaphase [27, 28]. The metaphase spindle must maintain a stable distance between the poles with chromosomes in the spindle mid-zone while maintaining an equal distance from each pole via dynamic microtubules. The astral MTs between the spindle pole and the cortex interact with the cell cortex through dynein motors and directly, inducing microtubule buckling. The dynein motors tend to elongate the spindle while the buckling force tries to compress it [9]. The metaphase spindle interzone hosts KMTs in connection with the kinetochores. The junction between them is occupied by minus end-directed motors (such as dynein), which in unison with KMT plus end depolymerization give rise to a contraction in the spindle [29]. The spindle contraction is countered by another set of proteins known as ‘chromokinesins’ operating between the KMT tips and the surrounding chromosomal arms. These exert pushing force on the spindle poles [30] tending to move them apart. The compressive and extensile forces compete to maintain the stable spindle configuration [9].

Spindle elongation occurs during anaphase and is one of the major events that characterize a successful mitotic division. It is the manifestation of increasing distance between the spindle poles that constitute the bipolar spindle. The anaphase onset triggers the breakage of cohesin spring at the KT-KT junction of sister chromatids, disrupting the inter-polar attraction force. This disturbs the balance of forces, allowing the segregation of duplicated chromosomes and the subsequent elongation and positioning of the spindle. Numerous experimental studies suggest that it is crucial for completing mitosis and forming healthy daughter cells. The segregation of duplicate chromosomes during anaphase is assisted [31, 32] by kinetochore-associated microtubule (KMT) dynamics (anaphase A) as well as elongation of the spindle (anaphase B). Thus, proper spindle elongation plays a role in ensuring that there is no chromosomal anomaly when the daughter cells are formed. It has been reported that in the blastomeres of *C. elegans*, the chromosome segregation takes place solely due to spindle elongation [33]. Timely elongation of the spindle positions the segregated chromosomes and centrosomes away from the center of the cell, determining the position of cytoplasmic cleavage during cytokinesis [8]. In this way, proper spindle elongation and positioning also facilitate cells to divide desired proportions of cytoplasm between the daughter cells.

Both the metaphase and anaphase spindles are mechanically robust against any external perturbations. In a recent experimental study conducted on *C. elegans* cells [34], optical tweezers were employed to displace one of the poles in a bipolar spindle, and the impact of this displacement was investigated upon release. The displaced pole returned to its original position, indicating that the spindle behaves like a spring. The repositioning of the spindle at the cell center was due to the dynamics of the astral microtubules generating strong pushing force on the poles. The stiffness of the spring-like force generator increases during anaphase and precisely position the spindle suppressing the thermal fluctuations [9, 34].

The mechanism involved in spindle elongation has been studied extensively in the past decades. Experimental studies on Drosophila embryos [35] revealed that spindle elongation involves non-KMTs present in the spindle interzone, called interpolar microtubules (IPMTs), which grow from one pole and interact with the IPMTs growing from the other pole in the spindle interzone. The *antiparallel* IPMTs (originating from different poles) have a finite overlap where they are cross-linked by mitotic motors like bipolar kinesin-5 [36, 37]. These motors can generate sliding motion on the IPMTs that in turn get exerted on the poles, which are thus slid apart [35]. Each IPMT has a free dynamic end in the spindle interzone and a less dynamic end anchored at the respective pole. The free MT end is the ‘plus end’ while the less dynamic end is the ‘minus end’. Before the onset of anaphase B, the IPMTs exhibit a net polymerization [35] at their plus ends, and net depolymerization at their minus ends. Because of this simultaneous polymerization and depolymerization of the IPMT plus ends and minus ends, respectively, the tubulin dimers that the IPMTs are composed of exhibit a net motion toward the respective poles. This rate of movement of the tubulin dimers along the length of the IPMT is referred to as IPMT flux rate [35]. The sliding action of the kinesin-5 motors on the IPMTs is antagonized by this IPMT flux rate of the tubulin dimers, preventing the spindle from elongating [35] under their sliding force. The mechanism ensures a constant interpolar separation.The onset of anaphase B reduces the rate of depolymerization at the IPMT minus ends, causing a decrease in the rate of IPMT flux [35, 38]. This reduces its antagonizing influence causing the action of kinesin-5 motors to become more effective in sliding the poles apart [35], thus resulting in the elongation of the spindle. Although this mechanism suffices in Drosophila embryos, it is not the general mechanism for spindle elongation. Experimental studies [39] have revealed that in most vertebrates, the astral MTs take part in spindle elongation. As stated earlier, the astral MTs emanate from each pole to grow and reach the cell cortex. The blastomeres of *C. elegans* showed marked dependence on these astral MTs for the elongation of the spindle [33] with the spindle interzone limiting the extent of this elongation. Most animals and protozoans depend on a secondary mechanism of spindle elongation [8] comprising of astral MTs. A study [18] in 2007 found that in the blastomeres of *C. elegans*, the kinesin-5 motor in the spindle midzone functions as a ‘brake’ on spindle elongation that resists the spindle’s extension in the face of a fast pole-separating mechanism. In such scenarios, the spindle elongation is assisted by the astral MTs [40].

Therefore, the IPMT mediated mechanism is limited by the IPMT overlap and needs a secondary mechanism to elongate the spindle. The astral MTs are found [32] to interact with the actin-rich cell cortex on the inner side of the plasma membrane to generate a motor-mediated pulling force on the respective poles, much like the way pulling forces are generated on the centrosome during Interphase. Astral MTs also buckle at the cell cortex due to their growth against the cortical barrier [21]. The growth generates a push on the poles directed away from the cell cortex toward the cell center. Therefore, besides the IPMT sliding force, the mechanism involves an interplay between the pulling and pushing forces [32] that ultimately translates into spindle elongation and its positioning, much like the way a single centrosome positioning occurs during Interphase.

The overlap in single centrosomal positioning and bipolar spindle elongation mechanisms has been utilized to craft a one-dimensional computational model that employs astral MTs and their interactions with the cell cortex. The single centrosomal dynamics employs the growth of the astral MTs from both sides of the centrosome, making their interactions the major force component. The bipolar spindle comprising of two individual poles employs similar tactics, but only on one side, between each pole and the proximal cell cortex. The MTs growing from each pole into the spindle interzone become the IPMTs and contribute to the sliding apart of the individual poles.

While the model for the IPMT-mediated spindle elongation has been studied earlier, it is limited by the overlap-dependent elongation [35]. As the poles separate, the IPMT overlap falls. The sliding motors operate with a sliding velocity close to their stall velocity, which is insufficient to attain the desired elongation or positioning of the spindle. Experimental studies [18, 40] indicate that in such circumstances, the astral MTs assist in spindle elongation. Although the mechanism is qualitatively explained, to our knowledge, a computational compilation of the same has not yet taken precedence. Here, we explore and develop upon an earlier model [35] for the IPMT-mediated elongation of the spindle and augment it with the astral MT mechanism as stated above.

While investigating the single centrosomal positioning in one dimension (1D), we exploit the basic model in the framework of a bipolar spindle for studying elongation kinetics of the latter. We further investigate these processes using elongated (elliptic) cells to include dynein-mediated sliding of the astral MTs along the cell cortex, which the 1D model cannot accommodate. The results from the 2D model with elliptic cells are consistent with 1D simulation. The models are implemented to explore the acceptable range of parameters that facilitate the desired patterns for single centrosome during interphase and bipolar spindles during mitotic anaphase B. The one-dimensional model can confer the duration of anaphase B and allow its application on rod-shaped cells like *C. elegans* [41] to a close approximation. The 1D model is further employed to study the impact of cell length (size), average MT length, the effect of MT buckling, and dynein-related force parameters on the centrosomal and bipolar spindle dynamics, while the extended 2D model is used to study the impact of differing cell shapes on the elongation kinetics of the anaphase B spindle, revealing exciting insights into their roles and applications.

## II. MODEL

### A. Positioning of a Centrosome

We propose a model for centrosome positioning in one dimension (1D). The forces acting on the centrosome (henceforth denoted by CS) in interphase cells are primarily mediated by astral MTs that grow to the cell cortex and interact with it. The minus-end directed cortical dynein motors bind to the astral MTs, causing them to exert a force on the CS toward the cell cortex. Again, astral MTs’ growth against the cell cortex causes them to buckle and apply a pushing force on the CS, directed away from the cell cortex. We assume that the astral MTs, modeled as polymeric rods, undergo first-order Euler buckling while growing against the cortex [22] generating a pushing force on the CS, which is inversely proportional to the square of the MT length. Thus, shorter MTs that reach the cell cortex generate a higher push on the CS. The dynein motors are uniformly distributed on the actin-rich cell cortex and quantified by a linear density. The number of dynein motors bound to a given astral MT in the cell cortex depends on the extent of the MT that grows beyond the cortex. A steric repulsion from the cell periphery confines the CS within the cell cytoplasm whenever an overlap is set to occur between them.

Fig. 1 shows the schematic of the mechanism of force balance. The region of interest on the cellular axis is identical to the cell length, which hereafter is referred to as the cell diameter and denoted by *d*_*cell*_. Thus, the cellular axis along the *X*-direction is bounded by the cell boundaries at *x* = 0 and *x* = *d*_*cell*_. The instantaneous position of the CS is denoted by *x*_*cent*_(*t*). The astral MTs undergo dynamic instability [42] in 1D, due to stochastic polymerization/de-polymerization, and are denoted by the position of their growing/shrinking tip *x*_*tip*_. The MT dynamic instability is modelled by a four-parameter system involving rescue frequency *f*_*res*_, catastrophe frequency *f*_*cat*_, growth velocity *v*_*g*_ and shrinkage velocity *v*_*s*_ [43]. Thus, for any given astral MT, we have:

**FIG. 1:**
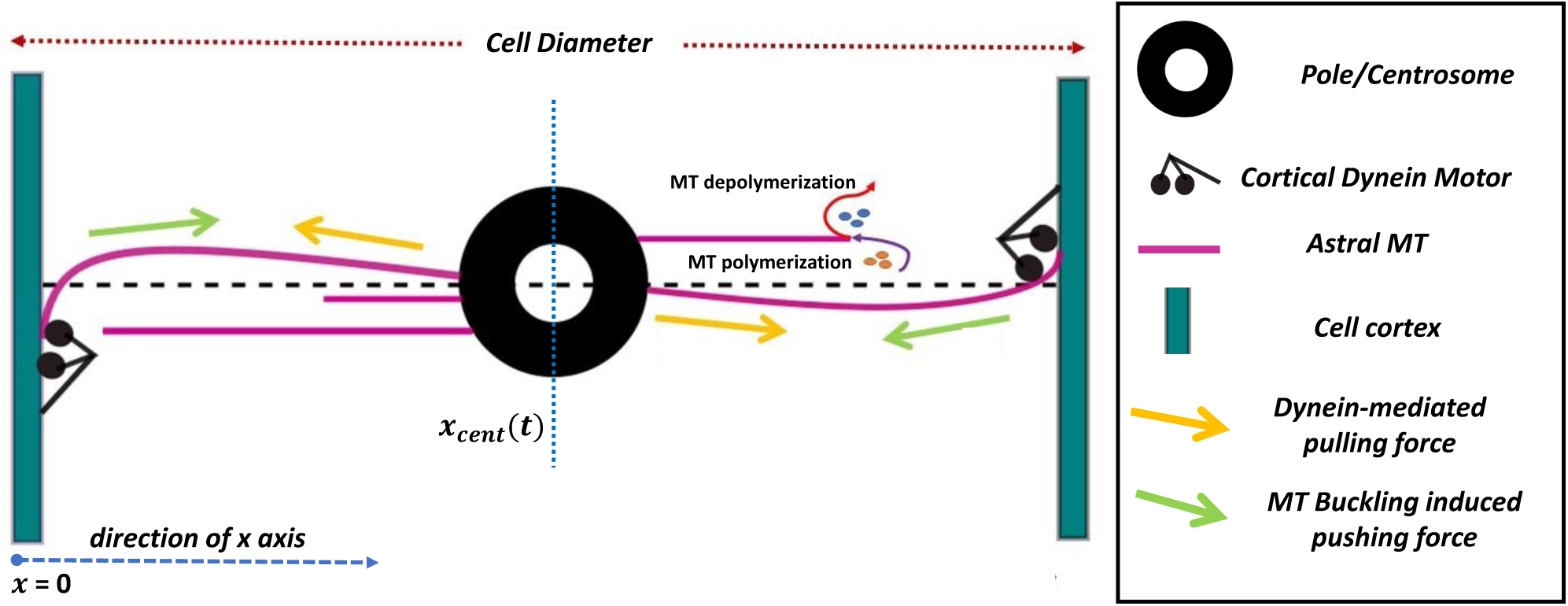
One-dimensional model for the positioning of a centrosome in a cell. The astral MTs undergo first-order Euler buckling upon growth against the cell cortex. The astral MTs exhibit dynamic instability indicated by depolymerization and polymerization of the MT tips. The arrows indicate the direction of the respective forces. The direction of the X-axis is shown in the picture. The horizontal dotted line, on which the centrosome rests, signifies the cellular axis along which it is restricted to move.

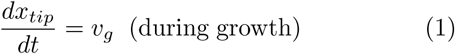

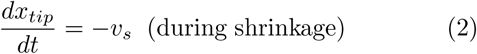

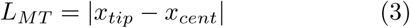

Where, *L*_*MT*_ represents the length of an astral MT emanating from the CS with it’s plus tip positioned at *x*_*tip*_. The average MT length corresponding to the dynamic instability parameters is given by [43, 44]:

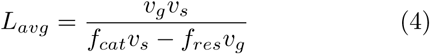

The amount of pushing force produced by an astral MT of unit length is defined by a buckling amplitude *A*_*buck*_ which appears to be about ten times the flexural rigidity of the astral MTs [11]. The linear density of dynein motors on the cortex is denoted by *k*_*dyn*_ and is uniform in character. Each motor engaged with an astral MT applies a mean force characterized by *F*_*dyn*_. MTs grow as per the dynamic instability parameters on both sides of the CS toward the cell cortex, where the position of the cortex is denoted by *x*_*cor*_.

The growth of the astral MTs against the cell cortex modifies the growth velocity [22] and the catastrophe frequency [45] causing the former to decrease and the latter to increase with progressing growth against the cell cortex. The growth against the cell cortex is not favored as it generates a pushing force opposite to the direction of growth of the MTs. The astral MTs exhibit resistance to this growth by modifying the dynamic instability parameters *v*_*g*_ and *f*_*cat*_ based on the pushing force experienced by them [12] as shown.

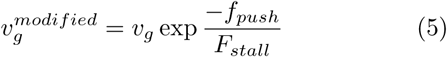

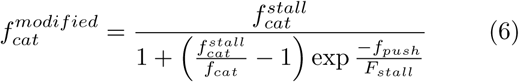

Here, *F*_*stall*_ refers to the load force on astral MTs that causes them to stall as they grow against the cell cortex; 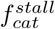 is the catastrophe frequency of a stalled astral MT [45] and *f*_*push*_ is the net load force that opposes MT growth against the cortex at a given instant. The modifications of the instability parameters stated in Eqs. (5) and (6) are applicable only after the astral MTs have reached the cortex. At the cortex, the MTs experience pulling and pushing forces. Let the pull experienced by the *i*^*th*^ astral MT emanating from the CS be 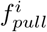 and the push be 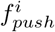. If 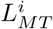 is the length of the i^*th*^ astral MT, we can write:

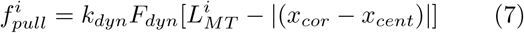

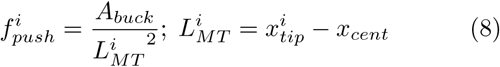

The length of an astral MT is measured from the position of its origin at the CS to its tip denoted by *x*_*tip*_. Subtracting the separation between the CS and the cortex from the MT length (see Eq. (7)), we obtain the length of the MT beyond the cell cortex. The dynein motors can bind to this length and contribute to pulling force. Summing over the superscript ‘i’ in Eqs. (7) and (8) would give the total pulling and pushing forces, respectively.

Now, suppose the CS moves toward the cortex with an instantaneous velocity. In that case, it experiences a drag force against its motion in the cell cytoplasm proportional to this velocity due to the drag coefficient of the cytoplasm, denoted by *μ*. Thus, the force balance equation for the CS becomes:

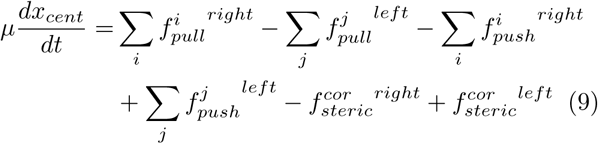

The superscripts ‘i’ and ‘j’ denote the astral MTs interacting with the right and left cortex, respectively. The steric repulsion force, denoted by 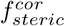, acts between the CS and the cell cortex when they almost overlap with each other and is inverse squared in nature. The cell cortex is separated from the cell boundary at *x* = 0 and *x* = *d*_*cell*_ by a distance of 0.75 *μm*. The steric force is quantified by:

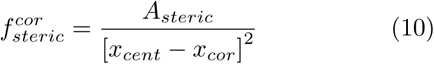

The amplitude of steric repulsion force is denoted by *A*_*steric*_. The buckling-mediated pushing and the dynein-mediated pulling forces apply only after the astral MTs have reached the cell cortex. Likewise, the steric force contributes to the repulsion between the cortex and the CS only after the CS has come within a distance of 1.5 *μm* from the cell cortex. We solve eq. 9 computationally to evaluate *x*_*cent*_(*t*) to obtain the stochastic dynamics of CS.

### B. Bipolar Spindle Elongation during Anaphase B

As shown in Fig. 2, the CSs or the individual poles in a bipolar spindle experience similar forces as discussed previously for the single CS, except in the spindle inter-zone. To describe forces in this region, we exploit the previous model of IPMT mediated spindle elongation in Drosophila embryo [35]. The spindle interzone consists of IPMTs interacting in antiparallel pairs (MTs emanating from opposite poles). Overlapping IPMTs get cross-linked by plus-end directed kinesin-5 that slide the MTs apart as shown in Fig. 2.

**FIG. 2:**
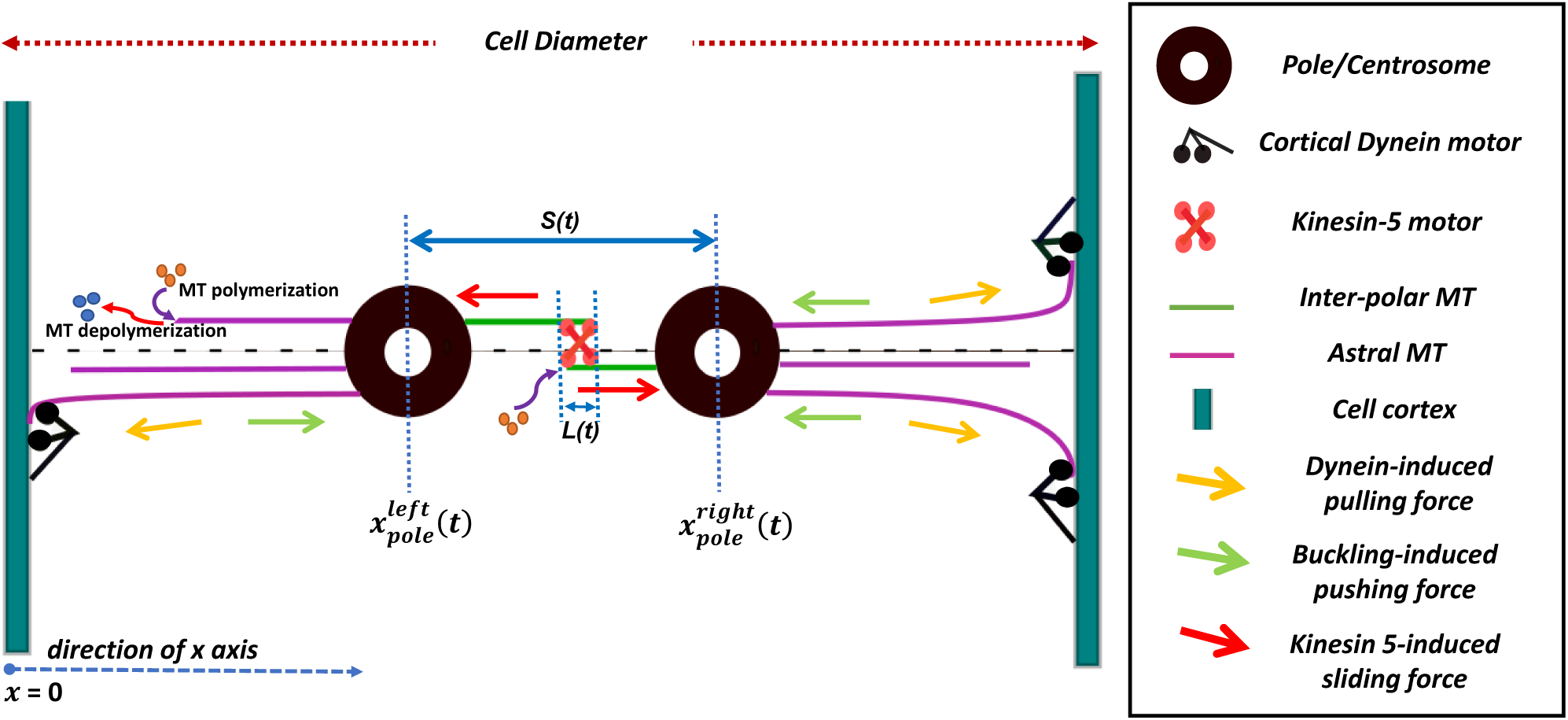
One-dimensional model for the elongation of bipolar spindle during Anaphase B. Here, the force-generating mechanisms mediated by the astral MTs on individual poles are similar to the single CS scenario as in Fig. 1. Antiparallel arrays of IPMTs crosslink in the spindle interzone via bipolar kinesin-5 which slide the IPMTs apart.

The IPMTs undergo de-polymerization at their minus ends and net polymerization at their plus ends [35]; meaning that tubulin dimers are deducted at their minus ends while there is a net addition of the same at their dynamic plus ends. The rate of de-polymerization of the IPMT minus ends corresponds to tubulin flux along the length of the IPMTs [35] and can be modeled via the poleward velocity of tubulin along the MT length, denoted by 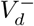. On the other hand, the rate of polymerization at the IPMT plus ends corresponds to the rate of increase of IPMT length and is denoted by velocity 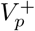. The IPMT flux of the tubulin dimers effectively antagonizes the sliding action of the kinesin-5 motors [35] present in the IPMT overlap zone, preventing the spindle elongation. The onset of anaphase B triggers a decrease in the rate of IPMT de-polymerization at the poles [35] causing the IPMT sliding to take effect. The kinesin-5 motor follows a linear force-velocity relation as shown below:

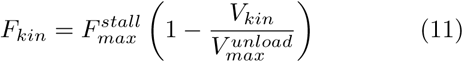

The kinesin-5 motor moves with a velocity *V*_*kin*_ under the load of *F*_*kin*_. 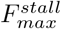 refers to the load under which the motor stalls causing the motor velocity *V*_*kin*_ to reduce to zero. Conversely, 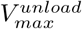 refers to the maximum *unloaded* velocity of the motor, that is, when the load force *F*_*kin*_ on it becomes zero.

The velocity of kinesin-5 equals the velocity with which the IPMTs are slid apart. Let the velocity be given by *V*_*sliding*_. The dynamic antiparallel IPMT overlap denoted by *L*(*t*), is shown in Fig. 2. Thus, we have [35]:

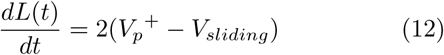

The factor 2 in the equation accounts for the assumed symmetry of the spindle interzone. Eq. (12) shows that the rate of overlap increases with increasing rates of IPMT plus-end polymerization and decreases as the rate of sliding of the antiparallel IPMTs increases. Now, let the instantaneous position of any one pole be denoted by *x*_*pole*_. Since the flux due to de-polymerization at the IPMT minus-ends opposes the pole separation, we have [35]:

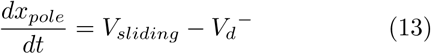

Factor 2 does not appear here because the above equation accounts for only one pole. The velocity of the pole’s movement, in Eq. (13), is favored by the IPMT sliding rates and is antagonized by increasing rates of tubulin flux along the IPMTs facilitated by the rate of IPMT minus-end depolymerization. As this pole moves through the cytoplasm toward the cortex, it experiences a drag force due to its drag coefficient *μ*, proportional to its instantaneous velocity. The resistance is countered by the action of the sliding motors in the spindle interzone. The number of kinesin-5 motors engaged between any set of antiparallel IPMTs is represented by a linear density *k*_*kin*_ and is proportional to the corresponding IPMT overlap. If *N* implies the total number of IPMT arrays in the spindle interzone [35], the force balance equation for any one pole is:

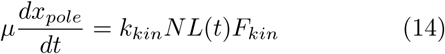

By algebraic manipulation of Eqs. (12), (13), and (14), we finally arrive at the following expression:

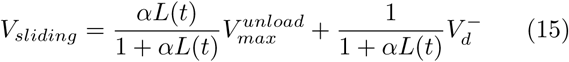

where 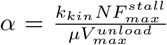 in Eq. (15) is a constant and denotes the strength of the IPMT sliding force against the viscous drag offered by the cytoplasm. Note that(see 15), for a small IPMT overlap, the sliding velocity tends to converge to the rate of de-polymerization at the IPMT minus end. Since this rate decreases at the onset of anaphase B, the elongation of the bipolar spindle is hindered [35]. Substituting Eq. (15) in Eq. (12) and (13), we obtain the IPMT contribution to the dynamics of any given pole:

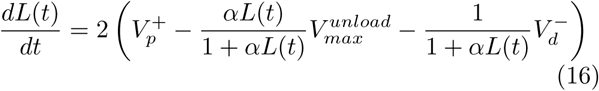

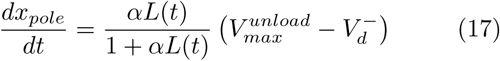

Since the spindle elongation is hindered due to decreasing overlap between the antiparallel IPMTs, the astral MTs must assist in further elongation. The force characteristic of astral MTs in a spindle is similar to the single CS, and the dynamic instability parameters of astral MTs emanating from one pole are independent of the other. The pulling and the pushing forces are given by Eqs. (7) and (8), respectively, whereas the altered dynamical parameters are given by Eq. (5), and (6). If the instantaneous position of the right and the left poles are denoted by 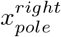 and 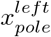 respectively, and length of the i^*th*^ astral MT is 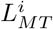, then we have:

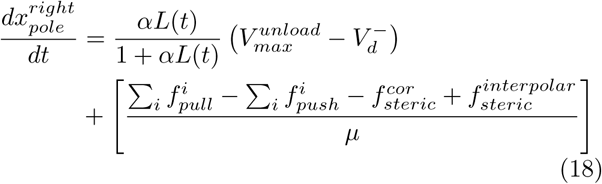

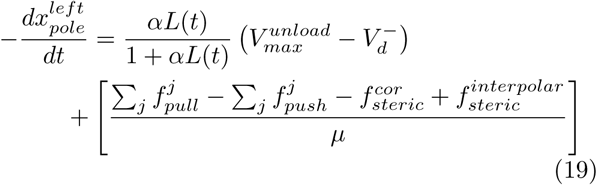

The ‘− ‘ sign on the left-hand side of Eq. (19) accounts for the position 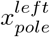 becoming smaller in magnitude as the spindle elongates and the left pole translates toward the left cortex. The superscripts ‘i’ and ‘j’ denote the astral MTs interacting with the right and left cortex respectively. The interpolar steric repulsion, denoted by 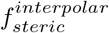, is included to avoid the poles from crossing each other:

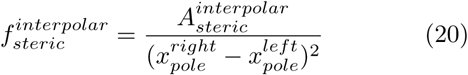

The amplitude of this steric force 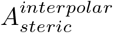 is non-zero only when poles are within a particular ‘cut-off’ proximity ∼ 1 *μm*. The steric repulsion between the cortex and the respective poles (i.e., 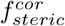) stays the same as in Eq. (10). Eqs. (16), (18) and (19) are simultaneously solved to obtain the interpolar separation *S*(*t*) as:

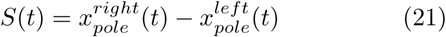

Thus, the model discussed here evaluates *S*(*t*) as the bipolar spindle elongation during anaphase B in a time-dependent manner.

### C. Modeling cortical sliding of astral MTs : an extension of the 1D models

The 1D models discussed restrict the astral MTs from growing along the 1D cellular axis. These MTs contribute to the net force in two crucial ways: MTs buckling at the cortex generate pushing force, and MTs binding to the cortical dynein motors generate pulling force. The interactions are microtubule length-dependent and occur in the space between the cortex and the cell periphery. The astral MTs being restricted to grow in one dimension, the contribution of these forces is limited by the cortical width. Restricted off-axis growth of the astral MTs at the cell cortex in the 1D model ignores the possibility of sliding along the cortex wall, preventing further interaction with dynein motors. Therefore, a considerable proportion of dynein motor-induced force is not considered by the 1D model. The 1D models are refined, accomodating lateral growth of the astral MTs along the cell cortex, and the contribution of cortical sliding is added to study the dynamics of the CS and the spindle.

In order to incorporate cortical sliding of the astral MTs, the cell geometry is updated from a 1D line to a long ellipse, providing the astral MTs freedom to grow and shrink in any direction in a 2D space. The CS (during Interphase) and the individual spindle poles (during Anaphase B) are still restricted to move along the 1D cellular axis (major axis) of the elliptic cell. This is to emphasize that the refined geometry is not a disjoint treatment but an extension of the previously discussed 1D models, incorporating additional complexity of a more realistic cellular environment.

#### 1. CS positioning during Interphase

Fig. 3(a) shows a schematic for positioning the CS during Interphase in an elliptic cell. The CS is constrained to move along the cellular axis (*x*-axis), represented by the ellipse’s major axis. The cellular axis is bounded by the elliptic cell periphery on both sides, at *x* = 0 and *x* = 2*a*_*x*_, where ‘*a*_*x*_’ is the semi-major axis of the ellipse. The cell’s transverse axis (*y*-axis) is given by ‘*a*_*y*_’. The astral MTs are allowed to emanate isotropically from the CS and have the freedom to undergo dynamic instability in 2D. The astral MT has a length *L*_*MT*_ and makes a polar angle *θ* (∈ [0, 2*π*]) with the *x*-axis, as shown in Fig. 3(a). Therefore, the randomly nucleated *i*^*th*^ astral MT has length 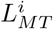 and polar angle *θ*^*i*^. Please note that,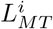 would follow the same dynamic equations for *x*_*tip*_ as in (1) and (2), but with *x*_*tip*_ replaced by 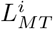 here. Corresponding to each astral MT, an angle *β*^*i*^ (see inset of Fig. 3(a)) is made by MT with the normal to the cell surface at the point of cortical contact.

**FIG. 3:**
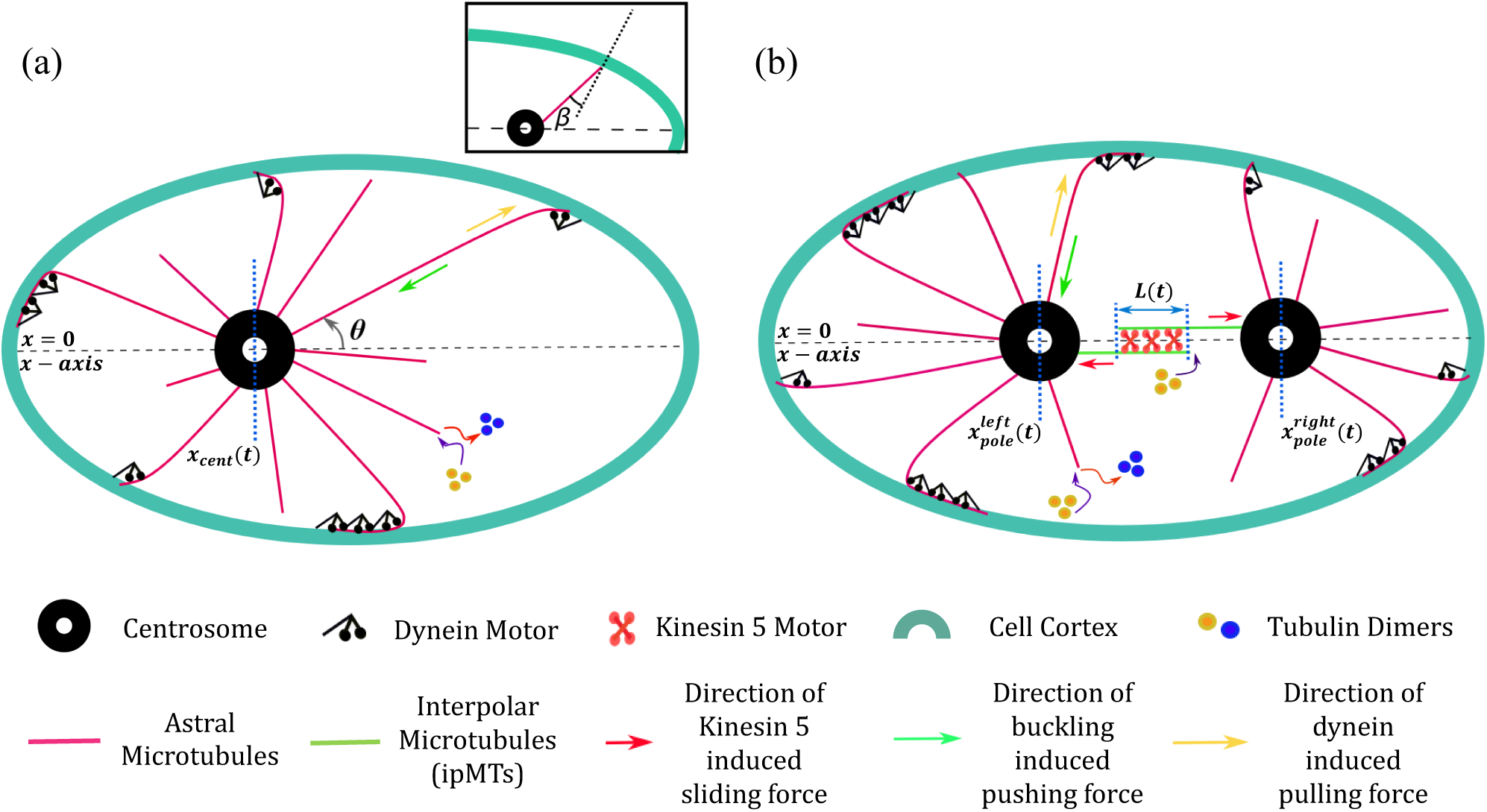
Models include cortical sliding of the astral MTs for positioning the interphase CS and elongation of the anaphase B spindle. (a) Interphase CS in an elliptic cell; the astral MTs emanate isotropically from the CS; the horizontal dashed line represents the cellular axis (x-axis) along which the CS is constrained to move; the orientation of each MT is denoted by the polar angle θ(∈ [0, 2π]); **inset**: β is the angle between an astral MT from the CS and the surface normal; the angle β determines the curvature-induced sliding (or catastrophe) with probability P_slide_; (b) model for elongation and positioning of the bipolar spindle during anaphase B in an elliptic cell; the IPMTs evolve along the cell axis, while the astral MTs evolve in 2D and can slide along the cortex; centrosomes/poles are constrained to move along the cell axis.

The astral MTs grow isotropically to the cell cortex and interact, applying forces on the CS. MTs buckle at the cortex while growing against it generating pushing a force 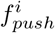 on the CS as in Eq. 8. The astral MTs bind to the dynein motors anchored to the cell cortex and generate a pulling force. The magnitude of this force depends on the extent to which the astral MTs grow within the cell cortex. In 1D model, they can grow up to the cell membrane (analogous to *β* = 0 in Fig. 3(a) with the MTs constrained to move along the *x* axis); therefore, the force exerted by dynein is strongly dependent on the thickness of the cortex. In the elliptical cell, the astral MTs do not always hit the cortex perpendicularly (i.e., *β* is not always equal to zero), due to the possibility of sliding along the cortical wall. Consequently, astral MTs can grow longer than the cortical width after reaching the cell cortex. This allows many cortical dynein motors to bind to and contribute to the pulling force. The astral MT sliding is governed by a probability ‘*P*_*slide*_’, and is a function of the angle *β*. For the *i*_*th*_ astral MT, making an angle *β*^*i*^ with the normal to the cortical surface, the sliding probability is given by:

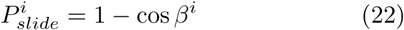

It is noteworthy to mention that since *β* is always zero in the 1D scenario, making the probability *P*_*slide*_ = 0. In the elliptical cell, a finite sliding probability (*P*_*slide*_ *>* 0) allows the astral MTs to slide along the cell cortex and contribute to the net force. Thus, the dynein motor induced pulling force generated via the *i*_*th*_ astral MT is:

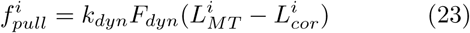

where, 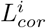 denotes the length at which the *i*^*t*^*h* MT hits the cortex as measured from its origin (CS), whereas 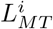 is the overall MT length. Thus, 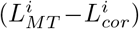 represents the segment of MT’s length sliding along the cortex. The parameters *k*_*dyn*_ and *F*_*dyn*_ have their usual meanings and values, as stated in Table I.

**Table I:** Parameters and their definition.

| *Parameter Definition (unit)* | *Parameter Abbreviation* | *Value* | *Range studied* | *Reference* |
| --- | --- | --- | --- | --- |
| Cell Radius ($\mu m$) | $r_{cell}$ | 20 | 10 – 30 | [11, 41] |
| Cell diameter ($\mu m$) | $d_{cell}$ | 40 | 20 – 60 | [11, 41] |
| Semi-major axis of the elliptic cell($\mu m$) | $a_x$ | 20 | 20,8 | This study |
| Semi-minor axis of the elliptic cell($\mu m$) | $a_y$ | 8 | 8,20 | This study |
| Steric Amplitude ($pN - \mu m^2$) | $A_{steric}$ | 150 | - | This study |
| Interpolar steric amplitude ($pN - \mu m^2$) | $A_{steric}^{interpolar}$ | 150 | - | This study |
| Astral MT stall force ($pN$) | $F_{stall}$ | 1.3 | - | This study, [22, 45, 46] |
| Astral MT average length ($\mu m$) | $L_{avg}^{AMT}$ | 40 | 5 – 60 | [11, 45] |
| Astral MT flexural rigidity($pN - \mu m^2$) | - | $\sim 18$ | - | [11] |
| Astral MT buckling amplitude ($pN - \mu m^2$) | $A_{buck}$ | 180 | - | [11, 45] |
| Number of astral MTs | - | 50 | - | [11] |
| Number of IPMTs in spindle interzone | N | 30 | - | [35, 47] |
| Drag Coefficient ($pN - sec/ \mu m$) | $\mu$ | 1000 | - | [48] |
| Stall force for Kinesin-5 ($pN$) | $F_{max}^{stall}$ | 1 | - | [49] |
| Unloaded Velocity of Kinesin-5 ($\mu m/sec$) | $V_{max}^{unload}$ | 0.07 | - | [35, 50] |
| Linear Density of Kinesin-5 on IPMT ($/ \mu m$) | $k_{kin}$ | 20 | - | [35, 51] |
| Force exerted by single cortical dynein motor ($pN$) | $F_{dyn}$ | 1.25 | - | [52, 53] |
| Linear Density of cortical dynein motor ($/ \mu m$) | $k_{dyn}$ | 5 | 1 – 5 | [45, 46] |
| IPMT catastrophe frequency ($s^{-1}$) | $f_{cat}^{IPMT}$ | 0.16 | - | [35, 54] |
| IPMT rescue frequency ($s^{-1}$) | $f_{res}^{IPMT}$ | 0.2 | - | [35, 54] |
| IPMT growth velocity ($\mu m/s$) | $v_g^{IPMT}$ | 0.16 | - | [35, 54] |
| IPMT shrinkage velocity ($\mu m/s$) | $v_s^{IPMT}$ | 0.16 | - | [35, 54] |
| Astral MT catastrophe frequency ($s^{-1}$) | $f_{cat}^{AMT}$ | 0.033 | - | [2] |
| Astral MT rescue frequency ($s^{-1}$) | $f_{res}^{AMT}$ | 0.067 | - | [2, 12] |
| Astral MT growth velocity ($\mu m/s$) | $v_g^{AMT}$ | 0.24 | 0.12 – 0.25 | [53, 55] |
| Astral MT shrinkage velocity ($\mu m/s$) | $v_s^{AMT}$ | 0.6 | 0.59 – 0.8 | [55] |
| Catastrophe frequency of stalled astral MT ($s^{-1}$) | $f_{cat}^{stall}$ | 0.042 | - | [12, 45] |

Pushing and pulling forces act along the length of the astral MTs, having an isotropic distribution with respect to the CS. Since the CS is constrained to move along the 1D cellular axis, only the component of force acting on the CS along the cell axis is essential. Given the orientation *θ* of each astral MT with the cell axis, we can obtain the projection of forces on the cell axis by multiplying them with cos *θ*. Let us refer to the model shown in Fig. 3(a). Due to the isotropic distribution of the astral MTs, angle *θ* falls in the range [0, 2*π*]. Note that pushing and pulling forces act along the length of an astral MT, directed towards and away from the CS, respectively. For astral MTs emanating from the right of the CS (or *θ* belonging to the first and the fourth quadrants as in Fig. 3(a)), the *x*-component of the buckling force (pushing) would cause the CS to move towards the left, while the dynein pull would cause the CS to move towards the right. Conversely, the pushing and pulling forces act in opposite directions for astral MTs emanating from the left of the CS (*θ* belonging to the second and the third quadrants).

At each time step Δ*t*, the net pushing and pulling forces acting on the CS are found by summing over the horizontal components of all such astral MTs that are interacting with the cortex. Therefore, at any time step, if the total pulling force toward the right side is 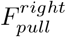 and that to the left is 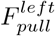, then we have:

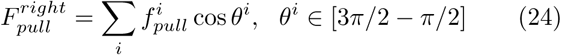

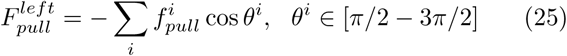

The −*ve* sign in Eq. 25 ensures that the sum of the force components evaluates to a positive value, as the cosine function is negative in the 2^*nd*^ and 3^*rd*^ quadrants. The index ‘*i*’ refers to all those astral MTs that are interacting with the cortex at a given time step. Similarly, if we denote the total pushing force towards the left and the right side as 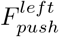 and 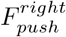 respectively, we have:

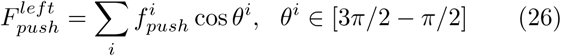

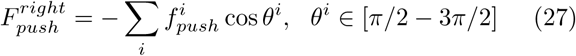

The forces given by Eqs. 24, 25, 26 and 27 that act on the CS at any given time, determine the dynamics of the CS during the interphase. Besides, a steric force from the cell boundary keeps the CS within the cellular confinement. If the instantaneous position of the CS along the cellular axis is denoted by *x*_*cent*_(*t*), the equation of motion can be written as:

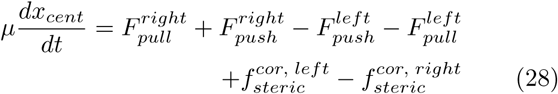

The steric forces on the CS at the left and right cortical boundaries have the same form, as mentioned in Eq. 10; ‘*μ*’ is the viscous drag coefficient of the cytoplasm. Numerically solving Eq. 28 yield the CS dynamics during the interphase, where the net force on the CS includes cortical sliding of the astral MTs, and forces applicable in the 1D model.

#### 2. Bipolar Spindle Elongation and Positioning during Anaphase B

Fig. 3(b) shows the model for the anaphase B spindle in an elliptic cell. The spindle has two sets of microtubules: IPMTs and astral MTs. The IPMTs’ interaction with the kinesin 5 motors is similar to the 1D model, shown in Fig. 2, whereas, the astral MTs are grown in 2D and can slide along the cortex. The spindle poles are constrained to move along the 1D cellular axis, represented by the long axis of the elliptic cell. Therefore, the *x*-component of the forces acting along the lengths of the astral MTs govern the dynamics of the individual poles. The sliding of the astral MTs at the cell cortex is dependent upon the angle *β*, which in turn determines the probability of sliding *P*_*slide*_, as per Eq. 22. *P*_*slide*_ dictates MT sliding at the cortex for each astral MT and determines the dynein-mediated sliding forces on the spindle poles. As discussed in the previous section, the evolution mechanics for the anaphase B spindle remains largely similar to the interphase CS in the elliptic cell, albeit with a few exceptions.

Each astral MT has a length *L*_*MT*_ and a polar angle *θ* that it makes with the cellular axis. The astral MTs do not emanate isotropically from each pole, unlike the single CS in the interphase cell. This is because the MTs existing between the poles are primarily of interpolar signature. The initial distribution of the astral MTs from each pole is restricted in the space between the pole and the proximal cell cortex. For the left pole, the initial distribution of the astral MTs at *t* = 0 *s* is restricted to the region left of the pole, in the 2^*nd*^ and the 3^*rd*^ quadrants. Conversely, for the right pole at *t* = 0 *s*, the initial astral MT distribution is constrained in the region right of the pole, in the 1^*st*^ and the 4^*th*^ quadrants. This condition applies to the poles only initially. As time progresses, the restriction is spontaneously removed, and the MTs can explore the cellular region, irrespective of their pole of origin. Thus, even if we do not have astral MTs directed towards the distal cell cortex initially, the evolving spindle eventually takes care of its contribution. Schematic in Fig. 3(b) shows an intermediate-time scenario, when the left pole has some astral MTs to its right (in the 1^*st*^ and the 4^*th*^ quadrants), and the right pole has some astral MTs to its left (in the 2^*nd*^ and the 3^*rd*^ quadrants), besides initially distributed arrays of MTs. As time progresses and the spindle undergoes elongation, each pole has astral MTs in all four quadrants. The situation for each pole is now similar to a single CS during interphase, as both pulling and pushing forces act on each of them, with their horizontal components directed towards the left and the right side.

The pushing and pulling forces that act along each MT are given by Eqs. 8 and 23. The horizontal components of these forces act on the poles along the cellular axis, with the contribution on each pole being independent of the other. To find the total horizontal pushing and pulling forces acting on each pole, their *x*-component are summed over at each time step, as shown in Eqs. 24, 25, 26 and 27. Therefore, for each pole, we have a total pulling force *F*_*pull*_ and a total pushing force *F*_*push*_ acting on them along the cellular axis, with directions to the right and left at each time step Δ*t*. Thus, we have:

i. *For θ*^*i*^ ∈ [3*π/*2 − *π/*2]

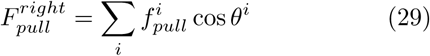

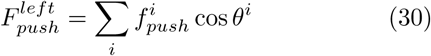
ii. *For θ*^*i*^ ∈ [*π/*2 − 3*π/*2]

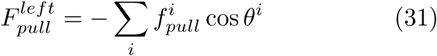

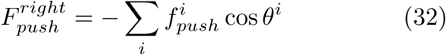

The -ve sign in Eqs. 31 and 32 appears due to cosine in the 2^*nd*^ and 3^*rd*^ quadrants. The index ‘*i*’ refers to the astral MTs that emanate from a given pole and interact with the cell cortex.

Since the IPMT dynamics remain unchanged from the previously discussed 1D model for the anaphase B spindle, its contribution to the net force has the same form as in Eqs. 18 and 19. Two steric forces act on each pole to confine the poles within the cell cytoplasm and prevent them from crossing each other. Designated as 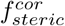 and 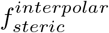 respectively, they have the same form and definition as in Eqs. 10 and 20. Therefore, if the instantaneous position of the right and left pole be denoted by 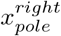 and 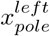 respectively, we invoke Eq. 16 and 17 and write the force balance equations for them follows:

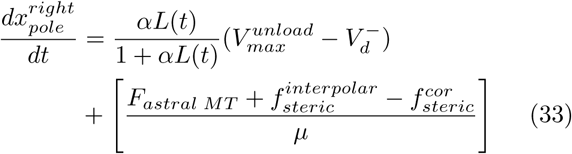

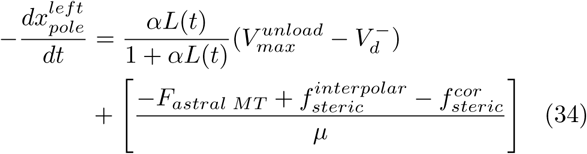

where,

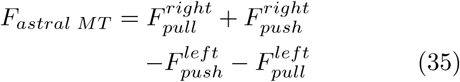

The ‘− ‘ sign on the left-hand side of Eq. (34) accounts for the position 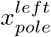 becoming smaller in magnitude as the spindle elongates and the left pole translates toward the left cortex.

Solving Eqs. 16, 33 and 34 simultaneously, we obtain the dynamics for the right and the left poles. We can obtain the spindle length at a given instant from the positions of the individual poles along the cellular axis using Eq. 21.

## III. COMPUTATIONAL TECHNIQUES

The MTs were assigned dynamic instability parameters (refer to Table I) exhibiting stochastic growth and shrinkage motion at their plus ends. In contrast, the minus end remained attached to the centrosomal poles. The dynamic instability at the MT plus ends was governed by the switching probability between events of growth and shrinkage during each time step Δ*t* (taken as 0.1 s), denoted by [56]:

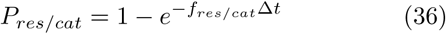

Here, *P*_*res*_ refers to the probability of the MT plus ends switching from shrinkage to growth and *P*_*cat*_ is the probability of the MT plus end switching from growth to shrinkage. *f*_*res*_ and *f*_*cat*_ are the frequencies corresponding to rescue and catastrophe events. If a random number called between 0 and 1 is found to be less than *P*_*res*_, the dynamic plus end switches to growth phase, and vice versa. This process is carried out at each time step using Eq. 36 to simulate the dynamic instability at the MT plus ends.

In the 1D model, the astral MTs that reaches the cell cortex, can grow till the cell periphery along the cellular axis. Upon hitting the cell boundary, MTs undergo catastrophe. In contrast, the astral MTs in the 2D elliptic cell have the option to slide along the cortex. Besides boundary-induced catastrophe, it can experience spontaneous catastrophe while executing dynamic instability at the plus end. MT sliding at the cell cortex is governed by the probability *P*_*slide*_ as in Eq. 22. A random number is called between 0 and 1 at each time step to determine the probability that the astral MTs would slide along the cortex or undergo catastrophe at the boundary.

For the single CS positioning, simulations were carried out for different initial positions of the CS, varied in steps of 5 *μm* along the cellular axis, including extreme and intermediate cellular zones. The astral MTs’ tip positions at *t* = 0 *s* were randomly distributed between the CS and the cell cortex on both sides. According to their dynamic instability parameters, the astral MTs grew in 1D to reach the cell cortex (see Table I). The random rescue and catastrophe events at their plus end allowed them to randomly interact with the cell cortex. A counter was introduced to estimate the number of astral MTs reaching the cell cortex at each step. Monitoring the dynamics of each astral MT in 1D(via *x*_*tip*_) allowed us to calculate their length *L*_*MT*_ (see 8). This data was used to evaluate the average dynein mediated pulling and MT buckling mediated pushing forces following Eqs. (7) and (8), acting on the CS at each time step. The pushing force so calculated was used to evaluate the modified growth velocity and the catastrophe frequency as per Eqs. (5) and (6). As indicated before, the MT growth on either side of the CS was dynamically independent. The counters measuring the number of astral MTs interacting with the cell cortex at any given time step were found to vary, implying fluctuations in their respective contributions to the resultant force. Using these data at each time step, Eq. (9) was solved to obtain the position of the CS.

To study the bipolar spindle elongation, the spindle was symmetrically positioned with an initial interpolar separation of 10 *μm* (held constant in all the simulations). The overlap of the IPMTs was obtained at each time step from Eq. (16) and fed into Eqs. (18) and (19) for obtaining the dynamics of the two poles. The astral MTs were simulated, similar to the single CS scenario. For IPMTs, only positive overlaps between the antiparallel arrays were considered. A negative overlap meant that the IPMTs were no longer overlapped. Thus, the sign of the overlap was checked at each time step. The stochastic dynamics of IPMTs allowed them to regain a non-zero overlap even if they had lost contact earlier.

The average polymerization rate 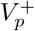 was found at each time step by monitoring the dynamic instability of the IPMTs and calculating their growth rate at each time step. 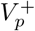 remained constant during each time step and was fed to Eq. (16) as a parameter. Also, since the depolymerization rate 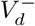 of the IPMT minus ends considerably decreases at the onset of anaphase B [35, 38], it was considered to be zero for simplicity. Using the expressions for the pulling and pushing forces from Eqs. (7) and (8), the forces were estimated for each pole. Based on the magnitude of the pushing force, dynamic instability parameters of the MTs were updated at each time step. Forces and IPMT overlap evaluated at each time step were used to solve the set of three differential Eqs (16), (18) and (19) simultaneously to obtain the interpolar separation as well as dynamics of individual poles.

In case of the elliptic cell, the astral MTs execute dynamic instability in 2D, with each assigned two parameters: an initial seeding length, and an orientation with the *x*-axis. The *x*-axis (cell axis) serves as the polar axis, along which the centrosomes are constrained to move. Initially, at *t* = 0 *s*, the astral MTs are distributed isotropically around the CS for the single CS scenario. In respect of the bipolar spindle, the initial distribution of the astral MTs is random in the 1^*st*^ and the 4^*th*^ quadrants for the right pole, and in the 2^*nd*^ and the 3^*rd*^ quadrants for the left pole. MTs grow and shrink in 2D according to their dynamic instability parameters, governed by the switching probabilities. Length of each MT, denoted by 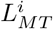 is updated at each time step Δ*t*. The probability of sliding *P*_*slide*_ is calculated for each astral MT at each time step to determine the cortical sliding and boundary-induced catastrophe once they reach the cell cortex. Instantaneous length of the astral MTs 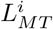 is compared with the respective length required for MT to reach the cortex. For all MTs having a length greater than that needed to reach the cell cortex, *x*-component of pushing and pulling forces acting on a given centrosome is estimated (see Eqs. 8, 23-27). Subsequently, Eq. 28 is numerically integrated to obtain the interphase CS position. Similarly, Eqs. 16, 33 and 34 are integrated for the right and the left pole simultaneously to obtain the kinetics of individual poles during anaphase B. The spindle separation is then obtained from Eq. 21.

All the above processes were repeated for 1000 ensembles to obtain the mean and the standard deviation in the data. The numerical codes were implemented in Fortran, and the differential equations governing the system dynamics were solved numerically using Runge-Kutta methods [57].

## IV. RESULTS

### A. Single CS Dynamics in 1D

#### 1. Buckling of astral MT plays a significant role in positioning the CS at the cell center

A single CS is considered in one dimension as a point on the cellular axis. The cell diameter is fixed at 40 *μm*. Fig. 1 shows the model schematic for studying the single centrosomal dynamics. As discussed before, the forces include astral MT buckling induced push (inwards, away from the cell cortex), cortical dynein motor mediated pull in 1D (outwards, toward the cell cortex), and a steric force to confine the CS within the cell cytoplasm. The model’s primary focus is to evaluate the interplay between these forces that dictate the force landscape of CS and bring about its positioning at the cell center. Fig. 4(a) shows centrosomal dynamics with initial position at *t* = 0 *s* of the CS at 5 *μm*, 15 *μm*, 20 *μm*, 25 *μm* and 35 *μm* along the cellular axis in a cell of diameter 40 *μm*. We observe in Fig. 4(a) that the CS stably positions itself at the cell center, irrespective of its initial position at *t* = 0 *s*. For initial positions at 5 *μm* and 35 *μm*, the CS is seen to move toward the cell center from the very beginning. As the CS is very close to the cell cortex at *t* = 0 *s*, short astral MTs emanating from it can easily reach the cortex. This interaction generates strong MT buckling mediated pushing force (since it is inversely proportional to the square of MT length acc. to (8)), causing the CS to move toward the cell center. The astral MTs from the CS on the other side (towards the distal cell cortex) have to grow to relatively longer lengths to reach the cell cortex. Thus, their contribution to the force landscape remains negligible at this point. As the CS’s motion progresses toward the cellular interior, the distance between the CS and the distal cell cortex reduces. As the distal cell cortex becomes more accessible to the astral MTs, their contribution to the force landscape of the CS gains significance. Since the net MT length remains comparatively long when the astral MTs reach the distal cell cortex, the buckling mediated force becomes less intense (refer to (8)), and their contribution to the force landscape thins out. As the CS moves toward the cell center and the distance between the CS and the distal cortex shrinks further, the buckling-induced force from the distal cell cortex becomes more prominent. The increasing buckling force from the distal cortex competes with the force contribution from the proximal cell cortex while maintaining a directed motion toward the cell center. At the cell center, the forces from the cell cortex on both sides of the CS become comparable. This eventually leads to attaining the force balance, thus stabilizing the CS at the cell center. The same also holds true for the CS initially placed at the cell center at *t* = 0 *s*. As seen in 4(a), the CS starting at 20 *μm* at *t* = 0 *s* is acted upon by the competing force landscapes from both sides that mediate its stay at the cell center. For the initial positions of the CS at 15 *μm* and 25 *μm* at *t* = 0 *s*, it moves toward the cell cortex for a period of ∼300 *s* at the beginning followed by movement toward the cell center. The initial placement of the CS at intermediate points along the cellular axis (15 *μm and* 25 *μm*) causes the astral MTs emanating from the CS to grow to greater lengths to reach the proximal cell cortex. This causes the buckling-induced force from the proximal wall to become less intense. The dynein-mediated contribution to the force landscape stays uncompromised as the astral MTs can ultimately reach the cell cortex and bind to the dynein motors. Therefore, the force landscape is dominated primarily by the dynein motor mediated pulling force that pulls the CS toward the proximal cell cortex during this initial period of ∼300 *s* as shown in Fig. 4(a). But this movement of the CS toward the cell cortex reduces the distance between the CS and the proximal wall, thus intensifying the buckling induced force. These strong inward forces acting on the CS then mediate its movement toward the cell center.

**FIG. 4:**
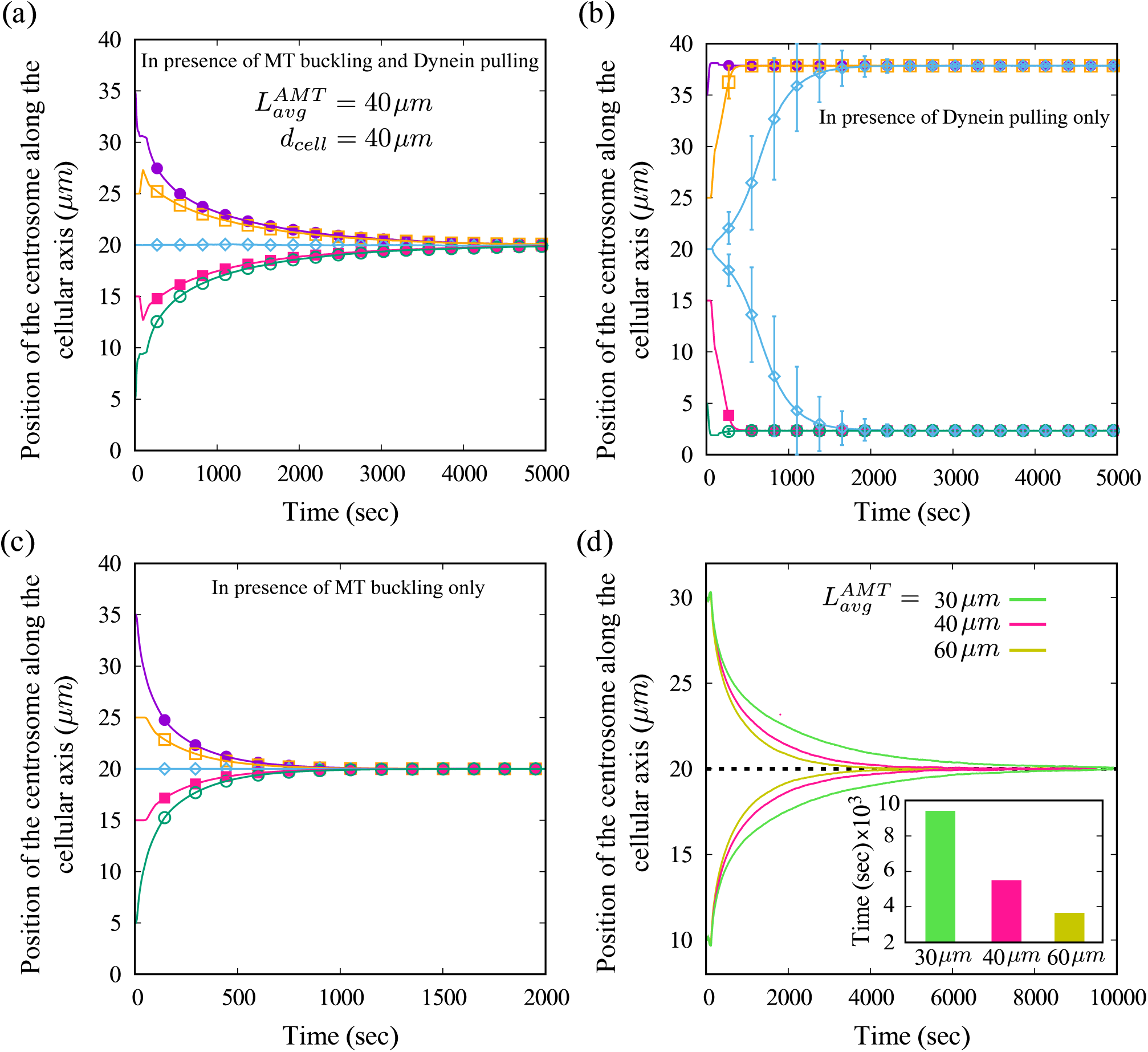
Variation in the position of CS over time is plotted for different initial positions at t = 0 s showing the role of astral MT buckling, cortical dynein motors, and average astral MT length. While figures (a), (b), (c) show CS dynamics for initial position at 5 μm, 15 μm, 20 μm, 25 μm and 35 μm at t = 0 s, figure (d) shows the same for CS initially placed at 10 μm and 30 μm. (a) The mean CS position along the cellular axis (in μm) is plotted versus time (in sec) for cell diameter 40 μm with astral MT avg. length 40 μm. (b) CS dynamics simulated without MT buckling (A_buck_=0); the two trajectories indicate two equally probable paths that the CS can take along the cellular axis with the given set of parameters. (c) CS dynamics simulated without cortical dynein (k_dyn_=0) showing that CS positioning at the cell center is dominated by force due to buckling of astral MTs. (d) The CS position plotted versus time for varying average MT length; **inset:** a bar graph showing duration required for CS positioning for various average MT lengths; faster positioning of the CS at the cell center is achieved by longer MTs.

To establish the roles of individual force contributors in positioning the CS at the cell center, we systematically exclude force contributors from our model one by one and examine their impact. We begin by setting the buckling amplitude *A*_*buck*_ to zero on both sides of the CS. In such a scenario, the CS does not experience any pushing force from either side of the cell cortex. The cortical dynein motors mediate the only force that acts on it. Fig. 4(b) shows the resultant centrosomal dynamics when buckling mediated forces are absent on both sides of the CS. For the CS initially placed anywhere along the cellular axis other than the cell center, the astral MTs reach the proximal cell cortex faster and more significantly in numbers than the distal cell cortex. This exerts a strong motor-mediated pulling force on the CS, directed toward the proximal cell cortex, away from the cell center. Thus, in the absence of buckling induced counter pushing force, the CS is dragged to the proximal cell periphery where the steric forces confine the CS.

For the CS placed at the cell center at *t* = 0 *s*, the astral MTs are equally likely to reach the cell cortex on either side of the CS. Thus, the CS has an equal probability of getting dragged to either the left or the right side when initially placed at the cell center. This is indicated in Fig. 4(b) by the two CS trajectories to the right and the left of the cell center. These two trajectories are equally probable and end up dragging the CS to the cell periphery in the absence of buckling mediated pushing forces, as shown in Fig. 4(b). Therefore, even though it may seem that the dynein mediated pulling forces from both sides of the CS should counter each other and achieve force balance, we find from Fig. 4(b) that dynein mediated forces do not exhibit such behavior, and are, therefore, not the main contributors for the stable positioning of the CS at the cell center.

We now proceed by excluding the force contribution of the cortical dynein motors on both sides of the CS such that no pulling force acts on it from either side. Fig. 4(c) shows centrosomal dynamics obtained by setting cortical dynein motor density to zero on both sides. The CS is seen to position itself at the cell center stably. Since only the buckling-induced force pushes the CS inwards toward the cell center, there is no counter force to oppose this movement due to the absence of cortical dynein motors on both sides of the CS. Thus, a CS starting at any point along the cellular axis except the cell center would be pushed toward the cell center under the net buckling force. The proximal cell cortex primarily mediates stronger buckling force than the distal cell cortex. But as the CS approaches the cell center, the buckling-induced force from the distal cortex intensifies and becomes more prominent. With the CS at the cell center, the buckling mediated pushing force from either side of the CS can compete with each other. Fig. 4(c) shows that this eventually leads to a force balance at the cell center that stabilizes the CS there. Thus, it is evident that astral MT buckling can position the CS at the cell center without the dynein motors’ assistance, as shown in Fig. 4(c). Therefore, we conclude that the astral MT buckling induced force is responsible for positioning the CS at the cell center during Interphase.

#### 2. CS rapidly positions at the cell center with longer astral MTs

In this section, we change the astral MT average length 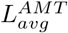 keeping the cell diameter constant to examine its impact on CS positioning. We start by altering the dynamic instability parameters of the astral MTs as they determine the 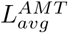 by eq. (4). We vary their velocities of growth and shrinkage while keeping their frequencies of catastrophe and rescue fixed (shown in Table I). The 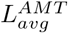 was varied with respect to the cell diameter *d*_*cell*_ by keeping it at 30 *μm* (*< d*_*cell*_), 40 *μm* (= *d*_*cell*_) and 60 *μm* (*> d*_*cell*_). Fig. 4(d) showcases the centrosomal dynamics obtained when the CS is initially placed at 10 *μm* and 30 *μm* along the cellular axis for varying 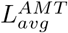. The results indicate that the time required for the CS to position at the cell center decreases when astral MTs present in the system have longer 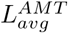. Since the centrosomal dynamics is governed by forces originating from the astral MTs’ interaction with the cell cortex, a longer 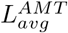 ensures stronger interaction between them, thus mediating faster positioning of the CS. The bar graph in the inset of Fig. 4(d) indicates the average time duration required by the CS to position at the cell center for differing astral MT average lengths. It explicitly shows that the positioning of the CS at the cell center is faster for astral MTs with a longer average MT length.

#### 3. CS position depends on the initial condition for short astral MTs

This section discusses the centrosomal positioning when astral MTs are relatively shorter with respect to the cell diameter. Fig. 5 showcases the results obtained from the 1D model for a cell of diameter 60 *μm* with 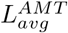 set at 40 *μm*. Fig. 5(a) shows that the centrosomal positioning does not occur at the cell center for these parameters. It is important to note here that astral MT buckling amplitude and linear density of cortical dynein motors, denoted by *A*_*buck*_ and *k*_*dyn*_, respectively, are kept unchanged as in the previous sections (see Table I). Fig. 5(a) shows the resultant CS dynamics for its initial placement at 5 *μm*, 25 *μm*, 30 *μm*, 35 *μm* and 55 *μm* along the 60 *μm* long cellular axis. The CS is placed very close to their respective proximal cell cortex for initial positions 5 *μm* and 55 *μm*. Therefore, many short astral MTs can easily reach the cell cortex generating strong buckling-induced forces. This causes the CS to initially move towards the cellular interior as shown in Fig. 5(a).

**FIG. 5:**
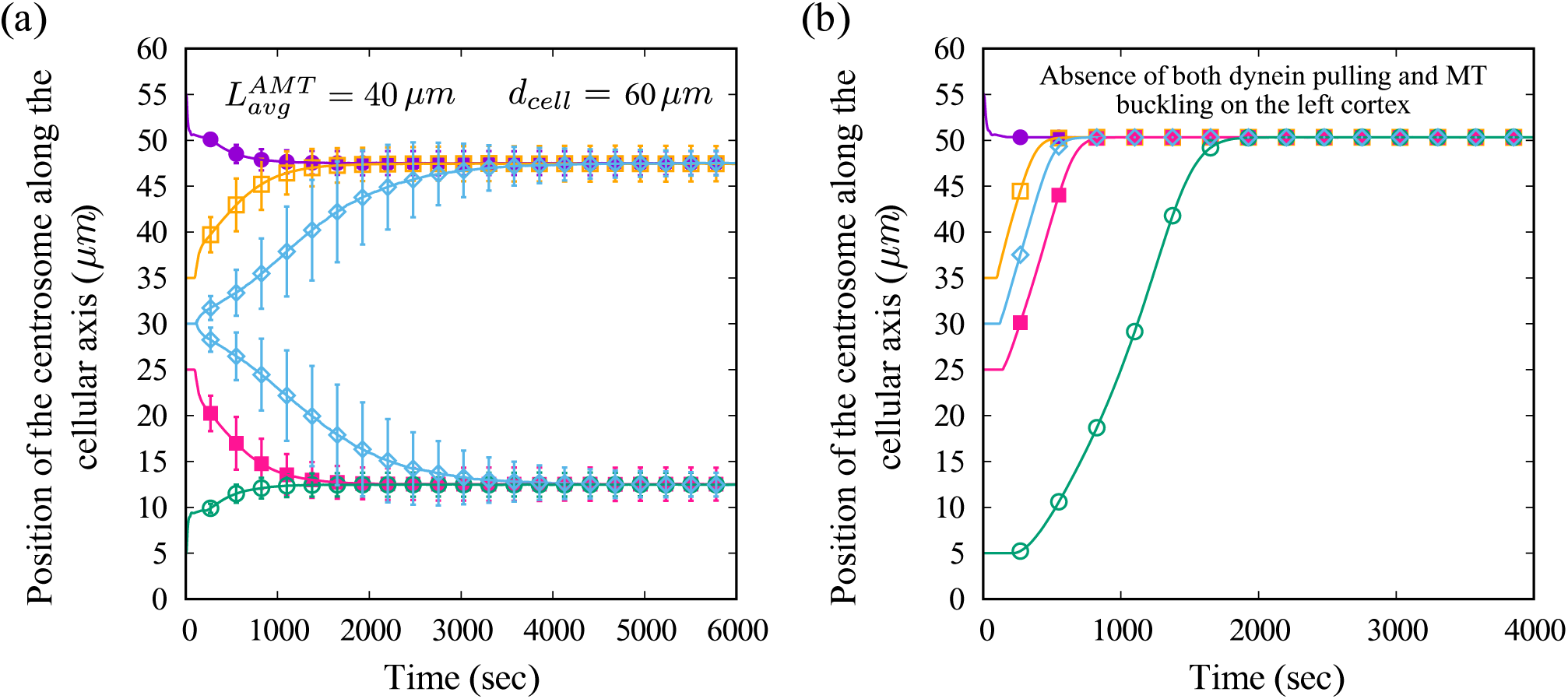
Dynamics of a single CS for short MTs with initial positions at 5 μm, 25 μm, 30 μm, 35 μm and 55 μm. (a) The mean position of the CS along the cellular axis plotted with standard deviation for cell diameter = 60 μm and avg. MT length = 40 μm; the two trajectories are equally likely paths that the CS can take along the cellular axis starting at the cell center. (b) Centrosomal position in absence of MT buckling and dynein pull from the left cortex (lower half of the plot) while the right cortex (upper half of the plot) is kept unmodified.

The movement of the CS towards the cell center increases the distance between the CS and the proximal cell cortex. The increased distance reduces the intensity of the buckling-mediated pushing forces allowing the dynein-mediated pulling forces from the proximal cell cortex to take precedence. These pulling forces pull the CS back towards the cell cortex, preventing its journey towards the cell center. The initial distance between the CS and the proximal cell cortex is greater for the CS starting at intermediate points along the cellular axis (at 25 *μm* and 35 *μm*). This allows the dynein-mediated pulling forces to play the dominant role during the initial movement of the CS. Thus, the CS is initially pulled towards the proximal cell cortex, as shown in Fig. 5(a). Again as the CS approaches the cortex, the buckling forces intensify, hindering its motion towards the cortex. Since the 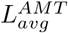 is short compared to the cell size, the force contribution from the distal cell cortex remains primarily insignificant. Therefore, the force landscape of the CS comprises an interplay between the pulling and the pushing forces mediated mainly by the proximal cell cortex. Eventually, a force balance results from this interplay, stabilizing the CS away from the cell center.

Exciting things happen when CS starts precisely at the cell center. In this case, there is an equal probability of the CS to position on either side of the cell center, shown by the two CS trajectories along the cellular axis towards the right and the left in Fig. 5(a). The astral MTs emanating from either side of the CS have to grow equal distances to reach the cell cortex on both sides. Since the cell size is large in comparison to the 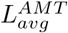 of the astral MTs, the length to which they must grow to reach the cell cortex on either side is long. This reduces the buckling-induced force, and thus, the cortical dynein motors dominate the net force landscape of the CS at the cell center. In the previous section from Fig. 4(b), we have concluded that the dynein-mediated pulling force cannot position the CS at the cell center. Therefore, the CS, under the influence of strong dynein pull, is dragged away from the cell center when the astral MTs start interacting with the cell cortex. Note that the astral MTs are equally likely to establish contact with the right or left cortex. Thus, the CS can position on either side of the cell center with equal probability, indicated by the equally likely CS trajectories shown in Fig. 5(a) when it starts at the cell center.

In Fig. 5(b), we tune the model such that the interaction between microtubule and cortex is weak on one side than the other. Such a possibility may arise for an elongated cell or when the MT array is asymmetric. In the model, forces are switched off on the left half of the cell (setting the buckling amplitude *A*_*buck*_ and dynein density *k*_*dyn*_ to zero) while keeping the right half undisturbed. The CS in Fig. 5(b), initially placed within the left half (at 5 *and* 25 *μm*), is seen to migrate toward the right half indicating that it is under the influence of an effective attraction originating from the right cortex. Moreover, CS stabilizes on the right half of the cell irrespective of the initial position. Thus, the force balance is independently established by pushing and pulling forces associated with the cell cortex on the right.

The dynamics of CS discussed in this context (Fig. 5(a)) can be modified by altering the force parameters. E.g., the buckling amplitude *A*_*buck*_, cortical dynein density *k*_*dyn*_, have a crucial effect on the CS position [discussed in Appendix I].

### B. Dynamics of Bipolar Spindle in 1D

#### 1. Astral and interpolar MTs together with cortical dynein robustly determine spindle kinetics during Anaphase B

During mitotic metaphase, the bipolar spindle poles maintain an approximately stable separation. The interpolar distance increases as the cell enters anaphase. A gradual slowdown follows this increase before a stable interpolar separation is achieved. The duration of the spindle elongation and subsequent positioning is observed to be in the range of 150-200 seconds in *C. elegans* [41] where the astral MTs play the pivotal role.

Fig. 6 shows the resultant spindle dynamics obtained from our model. Initially, the poles are placed symmetrically with respect to the cell center separated by a distance of 10*μm* (initial positions of individual poles are 20 *±* 5 *μm* on either side of the cell center) in a 40 *μm* cell. In Fig. 6(a), the variation in the interpolar distance is plotted versus time for two separate scenarios: with the (i) participation of IPMTs only and (ii) participation of both IPMTs and the astral MTs. The elongation of the spindle mediated by the IPMTs only, as shown in Fig. 6(a), continues to grow linearly and does not saturate. However, a closer look into the data indicates a faster separation in the early stages (see inset of Fig. 6(a) for interpolar separation in the first 20 sec). To address this, we investigate the IPMT overlap dynamics and the sliding velocity of the kinesin-5 motors that drive elongation of the spindle, as shown in Fig. 6(b). The crosslinking kinesin-5 motors maintain the overlap between the antiparallel IPMTs. In our model, the number of kinesin-5 motors actively participating in the elongation process is proportional to the IPMT overlap in the spindle interzone. The velocity of these motors along the IPMTs equals the rate at which IPMTs are being slid apart, given by *V*_*sliding*_. From the Fig. 6(b), we observe that the motor velocity *V*_*sliding*_ falls as the overlap *L*(*t*) falls, reaching a small value much less than their maximum unloaded velocity at 0.07 *μm/s*. This points to the kinesin-5 sliding motors operating near stall conditions as the IPMT overlap decreases. The phenomenon is reported earlier (15) where the motor velocity *V*_*sliding*_ is seen to depend on the dynamic overlap *L*(*t*) between the antiparallel IPMTs. The fall in the IPMT overlap thus accompanies decreasing sliding rates of kinesin-5 motors. As the IPMT sliding rate suffers a setback due to reducing overlap, the elongation of the spindle cannot proceed at the desired pace after the initial separation (∼5 sec). Also, since the IPMT overlap does not diminish entirely, as shown in Fig. 6(b), the sliding motors never actually stall completely, continuing the elongation process. Therefore, the mechanism mediated by IPMTs only helps push the poles apart but fails to give them a stable separation within the cell cytoplasm. In contrast, when the astral MTs participate alongside the IPMTs, the interpolar separation is seen to increase and then saturate as observed in the previous experimental studies [41].

**FIG. 6:**
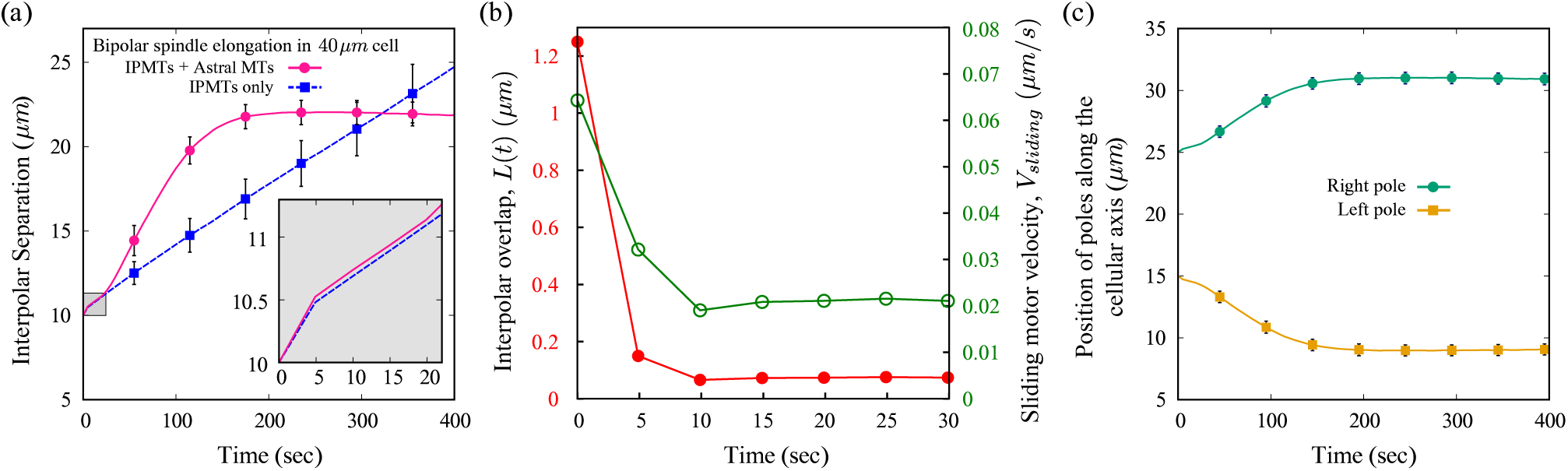
Bipolar spindle elongation during anaphase B. (a) The mean separation between the two poles constituting the bipolar spindle is shown for two cases when only the IPMTs participate and when the astral MTs and the IPMTs participate together. The cell diameter is kept at 40 μm with initial interpolar separation at 10 μm. The error bars signify the standard deviation. **Inset:** Bipolar spindle kinetics is shown for the first ∼20 s. The elongation up to this time frame is primarily mediated by IPMTs and is seen to decrease after the initial ∼5 s as an impact of decreasing antiparallel IPMT overlap. (b) The antiparallel IPMT overlap, L(t), is plotted versus time with the corresponding sliding velocity, V_sliding_, of kinesin-5, which drives the IPMT mediated bipolar elongation. (b) Mean position of individual poles is shown as a function of time when the IPMTs and the astral MTs participate together.

Upon comparing the elongation dynamics in Fig. 6(a) obtained for scenarios (i) and (ii) mentioned before, we find an initial overlap between them highlighted by the gray area. This spans the first ∼20 *s* when the spindle elongation is governed primarily by the IPMTs. Most astral MTs cannot reach the cell cortex during this period and contribute negligibly to the net force. Beyond this time frame, when the astral MTs actively participate, the poles are seen to elongate relatively faster than when IPMTs engage alone.

The astral MTs grow to reach the cell cortex and interact via MT buckling and cortical dynein motors binding to the astral MTs. The interpolar separation is small at the beginning of spindle elongation, while the distance between each pole and the proximal cortex is significant. The astral MTs emanating from each pole must grow long enough to reach the proximal cortex. As MTs reach the respective cortex, the corresponding buckling force (inversely proportional to the square of MT length (8)) starts to build up and push weakly on the respective individual centrosomes. At the same time, dynein-mediated pulling force (proportional to the cortical dynein density (7)) act on each centrosome and dominate over the buckling force. Therefore, the net force on the spindle imparted by the cortical dynein motors controls the initial elongation of the spindle resulting in a steady increase in the interpolar separation. As the separation between the poles increases, the distance between each pole and proximal cortex also decreases. Thus, many astral MTs can reach the cortex, and many of them are short in length. This leads to a change in the net force on the centrosome as the buckling force mediated by shorter astral MTs compensates the dynein-mediated pulling force. This hinders the elongation process of the spindle. As shown in Fig. 6(a), the elongation rates gradually decrease to zero, and the interpolar separation saturates. At this stage, a force-balance between dynein pull and MT buckling is achieved, preventing further elongation of the spindle. This is in contrast to the monotonic extension of the spindle without any saturation when only IPMT-mediated force contributes.

Fig. 6(c) shows the explicit movement of individual poles when both IPMTs and astral MTs participate. This movement corresponds to the net interpolar separation shown in Fig. 6(a). Note that, in this case, the poles were initially placed symmetrically about the cell center. The movement and final positioning of individual poles are also symmetric with respect to the cell center. The symmetry in spindle positioning is obtained even when the initial position of the spindle is not at the cell center (additional details are given in Appendix II).

#### 2. Astral MTs and cell size crucially regulate the rate of elongation and spindle length

Bipolar spindle elongation depends on astral MTs’ interactions with the cell cortex, which depends on the dynamic instability at their plus ends. The dynamic instability is determined by characteristic parameters *v*_*s*_, *v*_*g*_, *f*_*cat*_ *and f*_*res*_, which regulate the average length of MTs (see 4). In this section, we vary the average length of the astral MTs 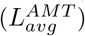 to explore its impact on the elongation and positioning of the bipolar spindle. The average length of the IPMTs is kept unchanged. Fig. 7(a) shows the spindle elongation dynamics obtained for 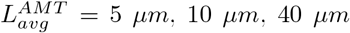 and 60 *μm* in a cell of diameter 40 *μm*. When 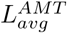 is less than the cell diameter (at 5 *μm and* 10 *μm*), the spindle elongation rates appear to depend on the 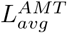 with shorter MTs taking longer to achieve a stable interpolar separation. For 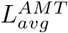 (40 *μm and* 60 *μm*) greater or equal to cell diameter; however, the elongation rates are fairly comparable. Steady-state position and extension of the spindle appear to be the same, irrespective of the variation in the average MT length.

**FIG. 7:**
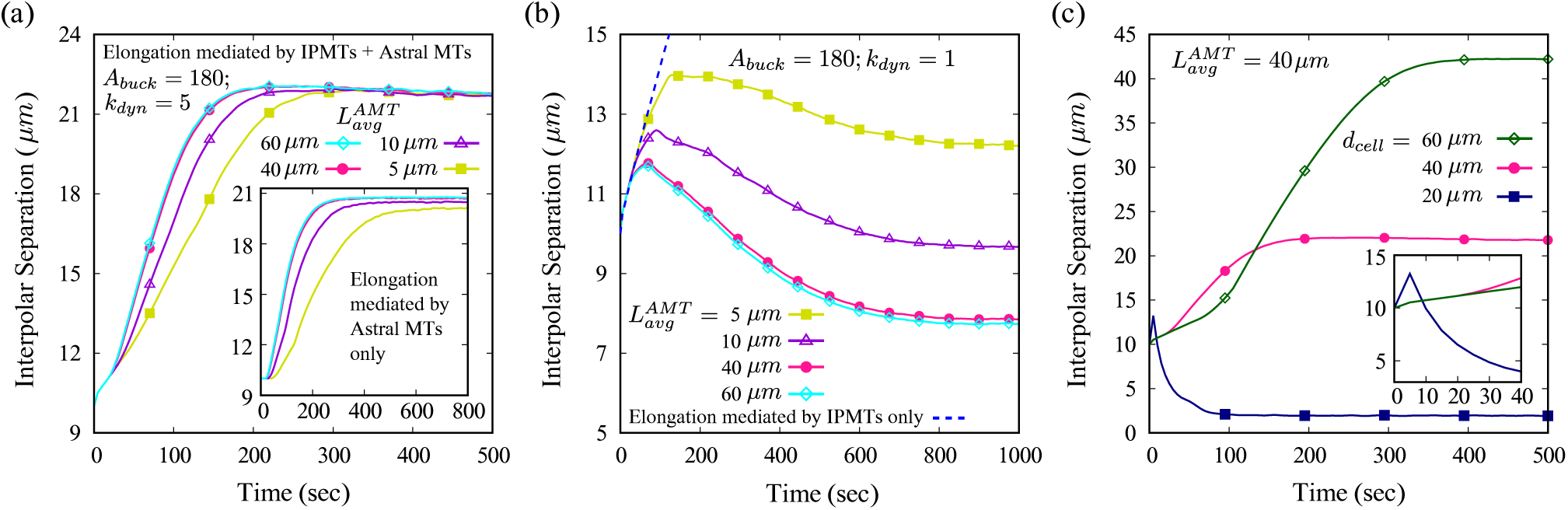
The impact of varying MT average length and cell diameters on bipolar spindle elongation. (a) The mean interpolar separation versus time shown for a 40 μm cell with average astral MT length of 5 μm, 10 μm, 40 μm and 60 μm; **Inset**: The mean separation obtained without IPMTs and only via astral MTs for the same set of MT length. (b) Interpolar separation similar to (a) but for small dynein density. (c) Kinetics of interpolar separation for varying cell diameters 20 μm, 40 μm and 60 μm, and average astral MT length 40 μm; **Inset:** Spindle dynamics shown during the first 40 s for differing cell diameters (as in (c)).

Notice that the spindle length is small in the early stage of the pole separation, and the distance between the poles and their respective proximal cortex is large. Thus, the astral MTs from a pole must grow long to reach the cell cortex. Long astral MTs cannot generate strong buckling force (see (8)), causing the net force on each centrosome to be dominated by dynein pull from the cortex. This causes a steady elongation of the spindle beyond the initial ∼20 *s* when the IPMT-sliding primarily mediates elongation (see Fig. 7(a)). Now, for a given separation between a pole and the proximal cortex, longer astral MTs reach the cell cortex faster than shorter MTs. This causes the spindle to elongate at a faster rate when 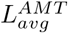 is greater. Hence, the spindle’s elongation rate varies with average MT length, as shown in Fig. 7(a). For astral MTs with an average MT length greater or equal to the cell diameter, the elongation rates almost coincide as the force contribution from the long astral MTs are very similar. As the spindle elongates, the distance between the poles and their respective proximal cortex shrinks. This allows shorter astral MTs to reach the cortex, increasing the buckling-induced force on the spindle, thus hindering the steady elongation of the spindle. With increasing buckling force, the net force on the spindle is no longer dominated by the cortical dynein motors. Spindle elongates gradually and then saturates when the pushing and pulling forces balance each other. In the inset of Fig. 7(a), the spindle elongation dynamics for different 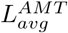 is plotted without IPMT force. With only astral MTs present in our model, we observe that the extent to which the spindle elongates depends on the average length of the astral MTs. The final interpolar separation almost coincides for 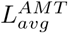 greater or equal to the cell diameter. For small values of 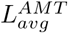, the final interpolar separation is relatively smaller and depends on the magnitude of 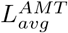. The result in the inset of Fig. 7(a) when compared to the main plot in Fig. 7(a), reveals that IPMTs not only determine the final spindle length but also keep it relatively independent of the average length of the astral MTs.

In Fig. 7(b), the spindle elongation obtained for various 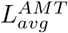 is plotted for a small dynein motor density on the cell cortex. The dotted line in Fig. 7(b) is the reference of spindle elongation when only IPMTs are present. The solid lines with points are for varying 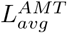 and consider both IPMTs and the astral MTs. The interpolar separation shows that the spindles initially elongate consistently, followed by contraction. The astral MTs, in contact with the cortex, generate both buckling and dynein-induced force. The buckling force becomes more influential here since the dynein density is low. Consequently, the buckling force dominates and pushes the poles away from the cortex, reducing interpolar separation. From this position, the astral MTs with higher 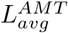 can reach the cortex faster and thus, buckle earlier. Therefore, the resultant spindle length with higher 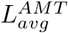 is smaller when dynein is impaired in the cortex.

We further explore the spindle elongation dynamics by varying the cell diameter while keeping the 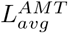 constant at 40 *μm*. The cell diameter is varied in steps of 20*μm* starting from 20 *μm* till 60 *μm*. The spindle evolution in Fig. 7(c) for cell diameters 40 *μm* and 60 *μm* shows similar elongation patterns that have been discussed so far. The primary IPMT mediated elongation continues for a longer duration in the 60 *μm* cell than the 40 *μm* cell. Also, the spindle elongation occurs to a much larger extent in the former, before the saturation sets in. This is due to the astral MTs that take longer to reach the cell cortex in a larger cell (with diameter 60 *μm*), slowing down the initial pole separation. The same factor also delays the exertion of buckling mediated pushing force on the poles, causing the spindle to elongate further. As for the cell with diameter, 20 *μm*, the astral MTs (with 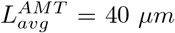) reach the cortex early and strongly pull on the poles outward, via cortical dynein motors. The cortical pulling causes the spindle to rapidly elongate in the beginning (see Fig. 7(c)) for the first ∼ 10 *s* (see inset of Fig. 7(c)). As the distance between the cortex and the poles shrinks, buckling force pushes the poles inward, hindering the steady elongation process. The cell size being small compared to 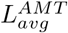, the buckling force becomes very strong after only a small elongation of the spindle, causing the poles to move toward each other steadily. In fact, for the smallest cell considered here (diameter 20 *μm*), the buckling force grows so intense that the spindle effectively collapses (see Fig. 7(c)).

#### 3. Bipolar spindle grows abnormally long in absence of astral MTs buckling

This section excludes the MT buckling force for studying the spindle separation kinetics. In the absence of buckling, IPMT sliding force pushes the poles apart. At the same time, cortical dynein motors pull the poles toward the cortex via astral MTs. The resulting spindle elongation dynamics with and without buckling force, shown in Fig. 8(a), indicate that the spindle separation is abnormally long in the absence of MT buckling force. As the poles approach near the cell cortex on their respective sides (see 8(b)), they encounter steric force from the cortex that restricts further spindle elongation. Thus, spindle length saturates close to the cell size (diameter). This is in sharp contrast to the typical spindle, as shown in Fig. 8(b). The spindle under normal conditions is relatively smaller, and poles are suitably positioned inside the cell. The buckling force thus plays a decisive role in placing the spindle poles within the cell cytoplasm and maintaining the spindle length.

**FIG. 8:**
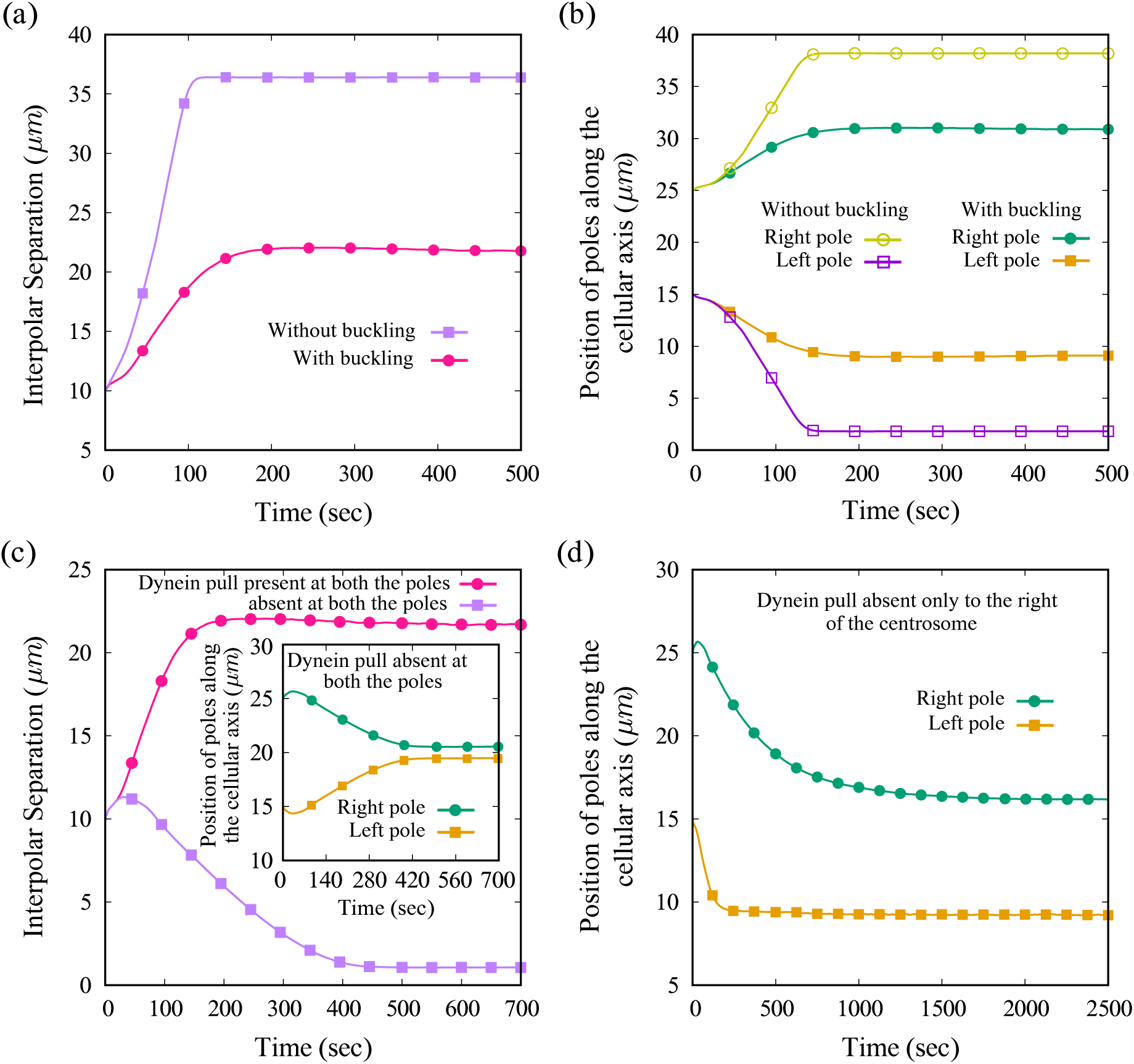
Bipolar spindle elongation and the role of dynein mediated pulling and buckling mediated pushing forces. (a) The mean interpolar separation is plotted versus time in the presence and absence of astral MTs mediated buckling. (b) The mean positions of the individual poles correspond to (a). (c) The interpolar separation is plotted versus time with and without dynein-mediated cortical pull via astral MTs. **Inset:** the mean positions of individual poles in the absence of dynein-mediated force on the poles. (d) The mean positions of the individual poles in the absence of dynein-mediated pull on the right pole.

#### 4. Bipolar spindle fails to elongate and position in absence of cortical dynein

This section excludes the dynein-mediated pulling force to examine their impact on the spindle elongation dynamics. The IPMTs slide and elongate the spindle under such a condition while the astral MT buckling force shortens the spindle. Fig. 8(c) shows the spindle dynamics in the presence and absence of dynein-mediated pulling force. The primary IPMT-mediated elongation can be seen in the beginning as the spindle elongates during the first ∼ 20 *s*. Once the astral MT buckling force acts on the poles, the elongation of the spindle is hindered. The inset in Fig. 8(c) shows the dynamics of individual poles with time. The poles move toward each other and tend to collapse, but the steric force between them avoids this possibility. Fig. 8(d) shows dynamics of individual poles when the dynein-mediated pulling force acts from one cortex only. Therefore, besides the IPMTs and the buckling forces as before, there is a contribution from the dynein-mediated pulling force on the spindle only from the left cortex (i.e., *y* ∼ 0 in Fig. 8(d)). The opposite cortex on the right (i.e., *y* ∼ 40 in Fig. 8(d)) has no cortical dynein motors and, therefore, offers no pulling force. In Fig. 8(d), we observe that the left pole can elongate and position normally, as found earlier. On the other hand, the right pole is pushed toward the cell interior due to buckling force from the right cortex. Finally, due to IPMTs sliding force, a force balance is achieved between the opposite poles that position the spindle asymmetrically about the center of the cell.

### C. Single CS and Bipolar Spindle Dynamics with cortical sliding of astral MTs in the extended 2D model

In this section, we present dynamics of the interphase CS and the anaphase B spindle using the mathematical model in 2D elliptic cell. The semi-minor axis is 8 *μm* long, and the semi-major axis is 20 *μm* matching the 1D cell diameter 40 *μm*. The centrosomes are constrained to move along the 1D cellular axis, shown in Fig. 9. Unless otherwise stated, all the other parameters remain unchanged (see Table I).

**FIG. 9:**
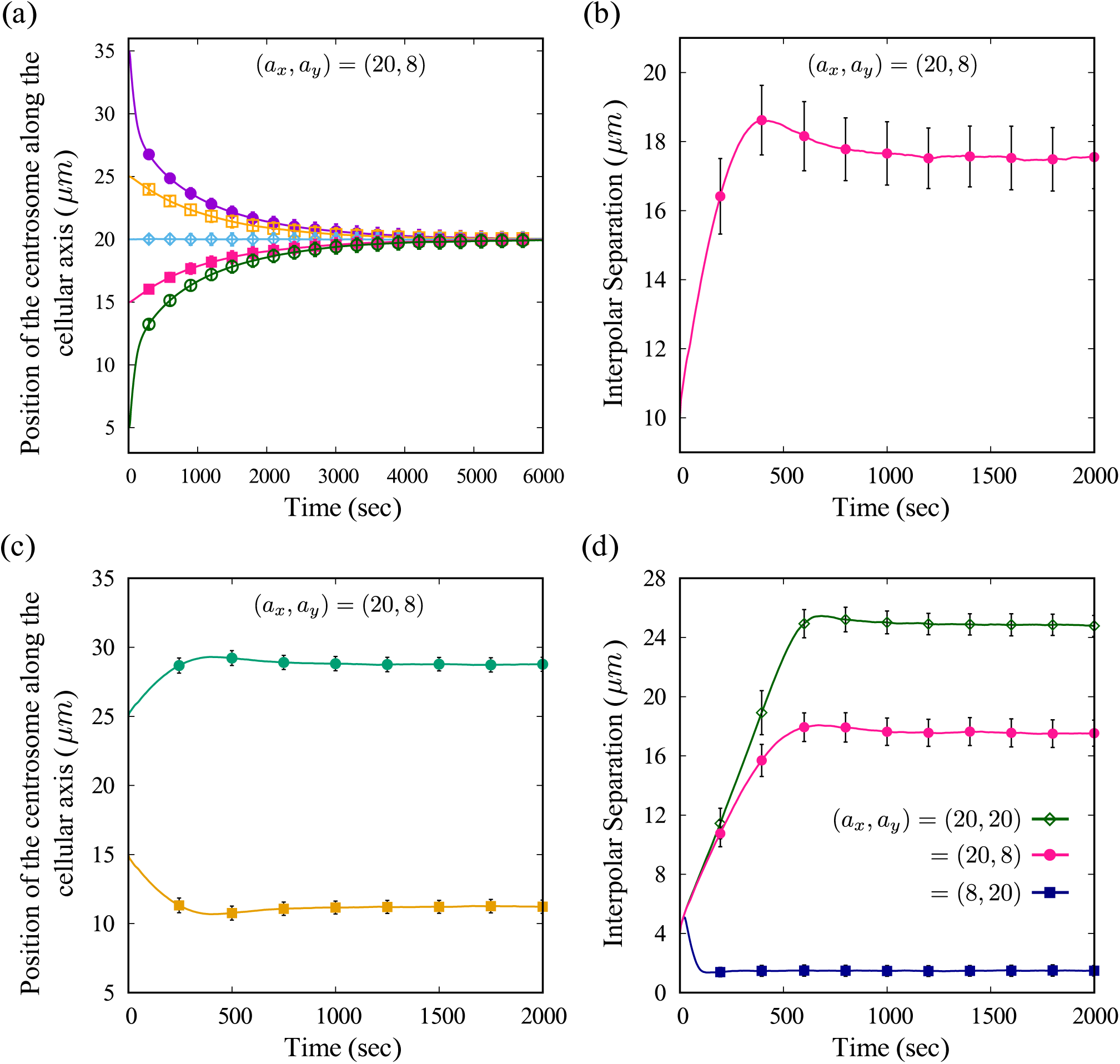
Dynamics of the CS during interphase and the spindle during anaphase B, as obtained from the model of elliptic cell with astral MT sliding at the cell cortex.(a): Simulation results showing positioning of the CS at the cell center during interphase;(b)spindle dynamics during anaphase B as obtained for an elliptic cell with semi-major axis a_x_ = 20 and semi-minor axis a_y_ = 8;(c) individual polar dynamics corresponding to the spindle dynamics shown in (b);(d) spindle dynamics as obtained for different cell shapes; circular cell(a_x_ = 20, a_y_ = 20),oblate cell(a_x_ = 20, a_y_ = 8;our standard elliptic cell discussed so far) and prolate cell(a_x_ = 8, a_y_ = 20).

Fig. 9(a) shows the CS dynamics during interphase as obtained from the model that includes astral MT sliding in a 2D elliptic cell. The CS can position itself at the cell center, as before, irrespective of the initial position along the cellular axis. Fig. 9(a) shows the dynamics of the CS starting close to the cell peripheries (5 and 35 *μm*) as well as intermediate regions (15 and 25 *μm*). Irrespective of the initial positions, the CS experiences strong pushing force away from the boundary, achieving a force balance at the cell center. Starting at the cell center (20 *μm*), the mean position of the CS continues to be the same as time progresses. Note that the force balance occurs at the cell center even with MT sliding added to the net force, thereby positioning the interphase CS at the cell center, as also observed previously in Fig.4(a) for our 1D model.

Fig. 9(b) shows the spindle dynamics obtained in the elliptic cell. Again we observe that the result obtained is qualitatively similar to the previously obtained results in Fig. 7(a). The spindle is observed to undergo elongation and subsequent positioning within the cytoplasm of the elliptic cell, much the same way it did for the 1D model. This is evident from Fig. 9(c), which shows the dynamics of individual poles coincide with the results shown in Fig. 7(c). The poles are observed to move steadily apart and then achieve a constant interpolar separation.

Fig. 9(d) shows the spindle dynamics obtained when the cell *shape* is altered. To alter the cell shape, we change the ratio between semi-major axis (*a*_*x*_) and semi-minor axis (*a*_*y*_). The plot in Fig. 9(d) shows the dynamics of the anaphase B spindle for three such cases: *a*_*x*_ *> a*_*y*_, *a*_*x*_ = 20, *a*_*y*_ = 8 (oblate), *a*_*x*_ = *a*_*y*_, *a*_*x*_ = 20 (circle) and *a*_*x*_ *< a*_*y*_, *a*_*x*_ = 8, *a*_*y*_ = 20 (prolate), with the cellular axis aligned along *a*_*x*_. The initial spindle length is kept at 4 *μm* for the three cases and all other parameters are kept unchanged.

Among these three cases, the spindle elongates to the greatest for the circular cell, followed by the oblate cell, while for the prolate cell, it largely fails to elongate, as shown in Fig. 9(d). The circular cell gives rise to longer astral MTs (since *a*_*y*_ = *a*_*x*_) in comparison to the oblate cell when they interact with the cortex, thus reducing the intensity of buckling force (since the buckling force is inversely proportional to the square of the astral MT length *L*_*MT*_ (see Eq. 8)). The oblate cell, on the other hand, gives rise to shorter astral MTs interacting with the cell cortex due to a shorter minor axis (*a*_*y*_ *< a*_*x*_). Thus, increasing the contribution of buckling force on the net spindle force. This allows the spindle to elongate more in the circular cell, in contrast to the oblate cell, as observed in Fig. 9(d). The prolate cell, on the other hand, has a short polar axis (directed along *a*_*x*_) along which the poles are now constrained to move. This greatly reduces the length of the astral MTs that may emanate from each pole toward the proximal cell cortex at small angles with respect to the *x*-axis. Such short astral MTs, upon interacting with the cell cortex, give rise to large buckling force (see Eq. 8). The force act along the length of the astral MTs, and make significant contributions to the net spindle force through their large *x*-component (horizontal) (see Eqs. 30 and 32). This effectively compresses the spindle by forcing the poles to move towards each other. Thus, the spindle fails to elongate when placed along the minor axis (along *a*_*x*_) of the prolate cell, as shown in Fig. 9(d).

## V. DISCUSSION

Positioning of the CS, like many other cellular organelles, is crucial for cellular functioning. While the CS position during Interphase is primarily dominated by MT buckling mediated pushing force, the bipolar spindle dynamics during Anaphase B is a consequence of the interplay between the buckling mediated pushing force and the dynein induced pulling, supported by the IPMT mediated sliding force. Our model for positioning of the single CS in 1D reveals that it is strongly driven by the cortical pushing force caused by the buckling of astral MTs interacting with the cortex. However, in the presence of short astral MTs or when the CS is placed in a relatively large cell, the mechanism allows for positioning of the CS away from the cell center. The stable positioning of the CS at the cell center can be facilitated by regulating the buckling amplitude of the astral microtubules and the cortical dynein density (see Appendix I). A mechanism similar to centrosome positioning is at work in fission yeast(*S. pombe*) cells during Interphase. The nucleus in *S. pombe* cells which is the microtubule-organizing center during Interphase, is positioned at the cell center by the microtubules emanating from its north and south poles. Experimental studies have shown that this mechanism is pushing force dominated [23]. Since our model reveals dominance of pushing force for centrosomal positioning in one dimension, the same model can also provide a qualitative understanding of the nuclear positioning at the cell center in cylindrical fission yeast cells during Interphase [23]. Even though the stable positioning of the single CS is influenced by MT buckling, the dynein density at the cortex can play a significant role in altering the CS position. Without dynein, the CS invariably positions at the cell center, and therefore altered positioning of the CS cannot be explained with MT buckling alone. We have also seen that the CS positioning is achieved at the cell center when the extended 2D model is represented by an elliptic cell. The cortical sliding of the astral MTs contributes to the pulling force, while the buckling at the cortex generates pushing force on the CS. Using the same set of parameters as in the 1D model, the net force position the CS at the cell center, indicating proper positioning of the CS during interphase. Therefore, a robust positioning of the CS can be achieved via MT buckling and dynein pull from the cortex.

Proper elongation and subsequent positioning of the bipolar spindle are crucial milestones for the cell to achieve faithful chromosomal segregation and cytokinesis during mitosis. The cortical dyneins are essential for spindle elongation. They help stretch the spindle by pulling force from the cortex, which is critical in anaphase B. However, a stable position of the metaphase spindle may be perturbed by a strong dynein pull from the cortex. Previous studies emphasized that cortical dynein pulling is anti-centering in nature, meaning that they tend to move the spindle away from the cell center [9, 34]. Yet, cortical dynein facilitates the force balance in unison with the antagonizing buckling mediated pushing force. The uniform MT-cortex interactions regulate a robust and symmetric spindle positioning in the cell mid-zone along the cellular axis.

The 2D model for the spindle in an elliptic cell also reveals that indeed, the dynein motor mediated pulling forces and the buckling induced pushing forces synergistically elongate and stabilize the spindle, even when the dynein-mediated pulling forces are enhanced by cortical sliding of the astral MTs. This mechanism is robust against changes in the cell geometry and cortical sliding of MTs. The kinetics of elongation and positioning of the anaphase B spindle in the elliptic cell is qualitatively similar to the 1D model. We further observe that the relative orientation of the spindle with respect to the cell axes may be significant during spindle elongation. When placed along the short axis of an elliptic cell, the spindle fails to elongate; however, the spindle elongates efficiently when oriented along the long axis.

A recent study by Farhadifar et al. [58] on nematode spindles has shown that cortical pulling forces are sufficient to position the spindle during anaphase B. Laser ablation experiments revealed that the pushing forces mediated by the cortex do not play a major role in mediating spindle’s positioning dynamics. Instead, it is the pulling force that significantly affects the positioning of the spindle, besides mediating the elongation. They emphasize the stoichiometric interaction between astral MTs and cortical force-generators(CFGs), the binding and unbinding rates of the CFGs, and the competition between astral MTs emanating from two different poles to bind the same CFG, as the main aspects of attaining the spindle positioning. Our models for the spindle focus on the overall impact of the force landscape mediated by astral MTs and the IPMTs on the spindle. It is possible to include the stoichiometric interaction (fixed number of CFGs per astral MT) between the astral MTs and the cortical dynein motors using our models. Nevertheless, in the absence of attachment and detachment of dynein motors, the modification of stoichiometric interaction in the present model may not be enough to reveal the pulling mediated positioning of the spindle. Our model for the single CS can invoke stoichiometric interaction between astral MTs and CFGs, and position the CS in an elliptic cell with cortical pulling forces alone (see Appendix III). Despite the fact that the bipolar spindle does not show a dynein pull-dominated spindle positioning, the single CS, in the presence of stoichiometric interactions, supports the cortical pulling-dominated positioning at the cell center.

The minimal models, considered here in 1D and 2D, have several limitations. For instance, uniform density of dynein on the cortex considered in the model cannot account for asymmetric cell division [19, 20] which could arise from the asymmetric positioning of the bipolar spindle (see Appendix II). Also, we employed a mean-field approach while evaluating the force exerted by each dynein motor and astral MTs interacting with the cortex. As mentioned before, additional features, including the pulling force-dominated positioning of the spindle, could be revealed if stochastic binding-unbinding of the cortical motors is considered. In addition, catch-bond-like binding of dynein motors [21] to the astral MTs with force-dependent attachment and detachment rates of the unbound and bound motors, respectively, may be another interesting aspect to consider in the future studies [59]. Another aspect ignored in this model is the absence of a force-generating mechanism involving the cytoplasm [9]. Experimental evidence accounts for cytoplasmic pulling influencing the centrosome’s movement, which may occur even when the astral MTs have not yet reached the cortex [14, 60, 61]. This pulling is possibly mediated by cytoplasmic dynein motors, either being anchored to actin filaments or by carrying vesicles along the microtubules [9, 14]. Our model does not incorporate cytoplasmic pulling force and thus cannot account for its impact on spindle elongation and positioning.

In our models for positioning the single CS during Interphase and bipolar spindle elongation during Anaphase B, we use the same set of parameters for studying two separate phenomena. We obtain stable positioning at the cell center for a single CS, while for two CSs of the bipolar spindle, we obtain faithful interpolar separation and positioning of individual poles. Even though we use the same parameters and the mechanism is largely similar for these scenarios, the single CS in 1D exploits the cortical interaction on both sides, while the individual poles in the bipolar spindle interact mainly with the proximal cortices. The microtubules from the opposite poles toward the cell center interact via Kinesin-5 motors, thus becoming the IPMTs. Under practical scenarios, this is feasible as the microtubules that grow in the spindle interzone mostly form IPMTs and KMTs. Thereby a tiny proportion of them become the astral MTs that can reach the distal cortex. Without the latter, the results still hold good; under practical circumstances, the number of such astral MTs is small and has no discernible effect on spindle dynamics. The elliptic cells require a more general distribution of the astral MTs such that the MT-cortex interactions are not limited only along the centrosomal axis. The interphase CS gives rise to an isotropic distribution of astral MTs, meaning that the cortical interactions are not just limited to the left and right of the CS. In the case of the spindle poles, the astral MTs emanating from each pole have an initial distribution that primarily interacts with the proximal cell cortex as in the 1D model; but eventually includes the distal cortex’s contribution as time progresses. Thus, the elliptic geometry facilitates a more realistic intracellular interaction.

An exciting feature of the model is that it can predict the spindle separation dynamics when MTs are short or the cell is large compared to the average MT length. As seen in the inset of Fig. 7(a), the final interpolar distance achieved during bipolar spindle elongation mediated by astral MTs alone with low 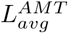 is relatively small compared to the usual scenario. The deficiency in pole separation dynamics is compensated by the IPMTs, which improve the extent of elongation and ensure that the final interpolar separation stays pretty similar even if shorter MTs mediated it. This provides the desired spindle elongation to facilitate proper cytokinesis.

Our model is based on a mean-field approach, and ignores molecular complexity. Despite that, the 1D model can account for the duration of bipolar spindle elongation in *C. elegans* [41] where the astral MT mediated mechanism dominates. Since we have characterized a cell in 1D by its diameter alone, our model can be mapped closely to rod-shaped cells like those of *C. elegans*. Irrespective of the parameters, qualitatively similar results are found for the interphase CS and the anaphase B spindle dynamics in 1D and 2D elliptic cells. Studying the impact of changing cell shapes on spindle dynamics provides valuable information about the spindle axis. With the correct choice of the spindle axis, the spindle can elongate during anaphase B determining the cell fate. Thus, despite a largely simplistic formulation, the models in 1D and 2D provide essential insights into an otherwise complex biochemical interaction.

Models also have limiting factors. The CSs are constrained to move along the x-axis, which serves as the centrosomal axis. Removing this constraint in 2D can reveal larger impacts of cell sizes and shapes on anaphase B spindles. Constraining the CSs along the *x*-axis restricts the plane of cytokinesis along the *y* −direction. Allowing the CSs to explore the 2D space freely would be useful to study variations in the cytokinesis plane during mitosis. A step further will be to model this system in three dimensions (3D) and examine the CS and the spindle dynamics in a more realistic space. This can further facilitate studying the impact of cell volume on spindle length and orientation and its scaling properties. Since both 1D and 2D models for the spindle are devoid of KMTs, the elongation kinetics of the spindle and chromosomal segregation ignore KMT dynamics [32, 62]. The models further do not account for interpolar attraction, mediated by minus end-directed motors in the spindle midzone, which can influence the spindle elongation during anaphase B [7, 63–65].

## VI. APPENDIX I

### PROPORTIONAL PULLING AND BUCKLING FORCES FROM CORTEX IS ESSENTIAL FOR CENTROSOME CENTERING WITH SHORT ASTRAL MTS

Earlier, we reported a single CS position away from the cell center with short astral MTs and a given set of interaction parameters, as shown in Fig. 5(a). This occurred because the force balance in either half of the cell was dominated by forces arising from the cortex within the same half, and contribution from the other half was insignificant. To achieve centering, net pushing force arising from both the cortex must be comparable. One possible way of achieving this is to increase the buckling amplitude while keeping the dynein density unchanged. This would facilitate more pushing force for the same microtubule length allowing the CS to counter dynein-mediated pulling force at larger distances from the cell cortex. Another way is to decrease the dynein density on the cortex so that the CS experiences less pull toward the cortex for the same distance from the cortex. Based on these hypotheses, we simulated the model with large buckling amplitude and dynein density (see Fig. 10(a)) and small buckling amplitude and dynein density (see 10(b)). In both cases, we found CS could position at the cell center, suggesting proportional buckling and pulling forces from the cell cortex is essential for CS centering with short astral MTs.

**FIG. 10:**
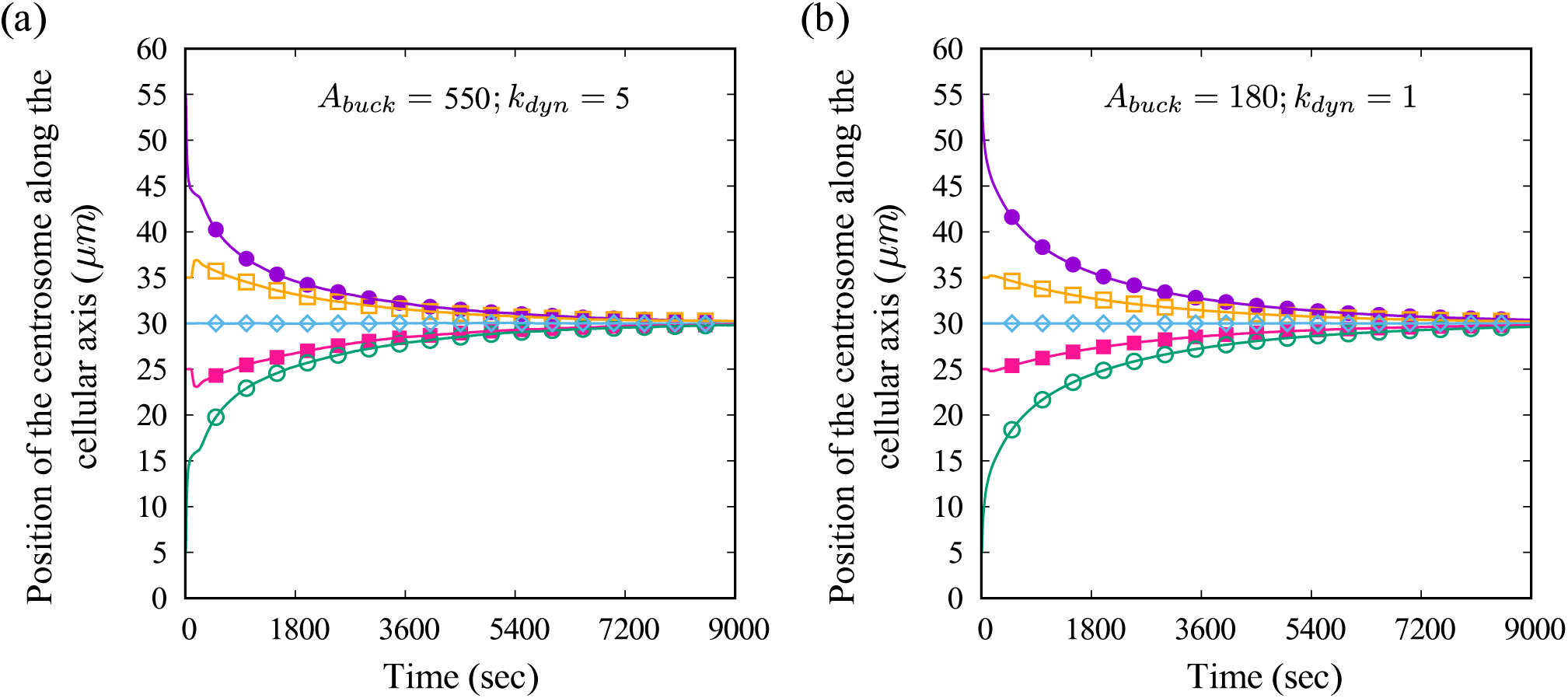
Single CS positioning in a 60 μm cell with short astral MTs (average length 40 μm), plotted for intial CS positions at 5 μm, 15 μm, 25 μm and 35 μm away from the left cortex. (a) The mean centrosomal position for large buckling amplitude and dynein density. (b) Mean centrosomal posiiton for small buckling amplitude and dynein density.

## VII. APPENDIX II

### SYMMETRIC POSITIONING OF THE BIPOLAR SPINDLE WITH INITIALLY ASYMMETRIC POLES

Fig. 11 shows dynamics of individual poles when placed *asymmetrically* at *t* = 0 *s* for normal and short astral MTs respectively. Asymmetric positioning of the poles refers to the initial spindle’s center shifted away from the cell center. In other words, one of the poles is closer to the cortex than the other at the onset of spindle elongation. In Fig. 11(a) we show spindle elongation dynamics of two sets of poles placed asymmetrically on either side of the cell. Data shows that the pole closer to the cell cortex moves relatively less and is positioned rapidly than the one away from the cortex. However, the final positioning of the spindle is symmetric with respect to the cell center. This occurs due to the uniform dynein density and astral MTs buckling on either side of the bipolar spindle. The spindle separation kinetics is slightly delayed with short astral MTs, as shown in Fig. 11(b). Nevertheless, the final spindle positioning and interpolar distance remain the same as observed with normal astral MTs.

**FIG. 11:**
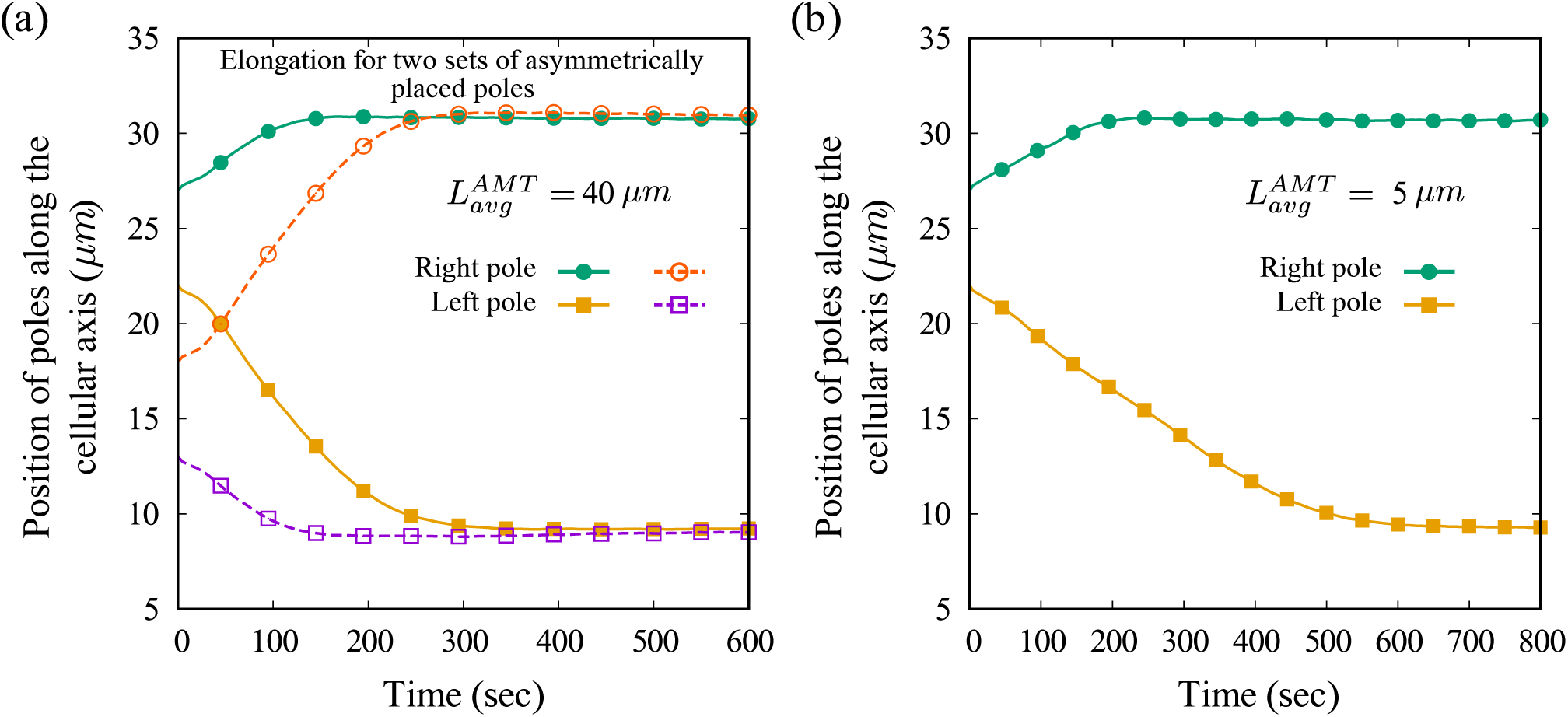
Symmetric positioning of poles that were asymmetrically placed at t = 0 s. (a) Final positioning and length of the spindle remain unaffected for varying initial positions of individual poles; (b) Pole separation kinetics for short astral MTs indicating that the symmetric positioning of the poles remain unaffected even when short astral MTs are present in the system.

## VIII. APPENDIX III

### SINGLE CS POSITIONING WITH STOICHIOMETRIC INTERACTION BETWEEN ASTRAL MTS AND CORTICAL DYNEIN MOTORS IN ABSENCE OF BUCKLING MEDIATED PUSHING FORCE

Stoichiometric interaction between astral MTs and cortical dynein motors was suggested in a recent study [58] to facilitate cortical pulling force that can dominate the positioning of CS in a cell. Here, we explore a similar scenario, including stoichiometric interaction between the astral MTs and the cortical motors for positioning a single CS in elliptic cells. Such interactions call for a fixed number of cortical force generators per astral MT. In our original model, we evaluated the cortical pulling force using Eq. 23, which depends on MT length and the linear density of dynein motors given by *k*_*dyn*_. We fix the number of dynein motors that each MT interacts with to ensure stoichiometric relation. The recent study [58] investigated this interaction using a 1 : 1 ratio, meaning that each MT interacts with a single cortical force generator (i.e., dynein motor). We explore the impact of a larger ratio of 1 : *k*_*dyn*_ (*k*_*dyn*_ = 5) on the CS dynamics, meaning that each MT has to interact with *k*_*dyn*_ number of motors. This causes the pulling force exerted by each MT evaluated by counting the number of motors *k*_*dyn*_ per MT where the force exerted by each motor is *F*_*dyn*_. The parameters are stated in Table I. Therefore, with stoichiometric interaction, the pulling force exerted by each astral MT interacting with the cortex remains the same, irrespective of its sliding length along the cortex. Therefore, even when the CS is placed near the cell cortex, the force on it from the proximal wall does not increase monotonically due to a limited number of dynein motors on each MT. As shown in Fig. 12, the CS, when placed on the cell axis at 5 *μm* and 35 *μm*, experiences a net pull toward the proximal cell cortex but eventually gets pulled toward the cell center due to the pull from the distal cell cortex. A similar outcome is observed when the CS is placed slightly off the cell center, at 15 *μm* and 25 *μm*. This points to the force contribution from the distal cortex dominating the CS trajectory until a force balance occurs at the cell center. Initially, the MTs emanating from the CS toward the proximal cell cortex are the primary force contributors that pull the CS towards the proximal cortex (Fig.12). When the CS advances towards the proximal cell cortex, more MTs emanating from it and interacting with the proximal wall become obliquely aligned with the centrosomal axis. This is because the astral MTs that interact with the cortex remain anchored at the cortex while the CS advances toward the proximal wall along the cellular axis. As a consequence of increasing oblique astral MTs, the magnitude of the horizontal force components contributing to the pull on the CS starts decreasing. At the same time, MTs emanating from the CS toward the distal cortex greatly align with the *x*-axis, increasing horizontal pulling force toward the distal cortex. Once the pulling force acting on the CS from the distal cell cortex becomes stronger than the proximal cortex, CS moves toward the cell center, as shown in Fig. 12. This eventually leads to a force balance, stabilizing the CS at the cell center. For the CS starting at the 20 *μm* mark, the horizontal components of the pulling forces coming from the opposite cortex are comparable, localizing the CS at the cell center. Therefore, our model for the CS positioning allows for dynein-mediated positioning of the CS at the cell center, when a stoichiometric ratio fixes the number of cortical motors per MT.

**FIG. 12:**
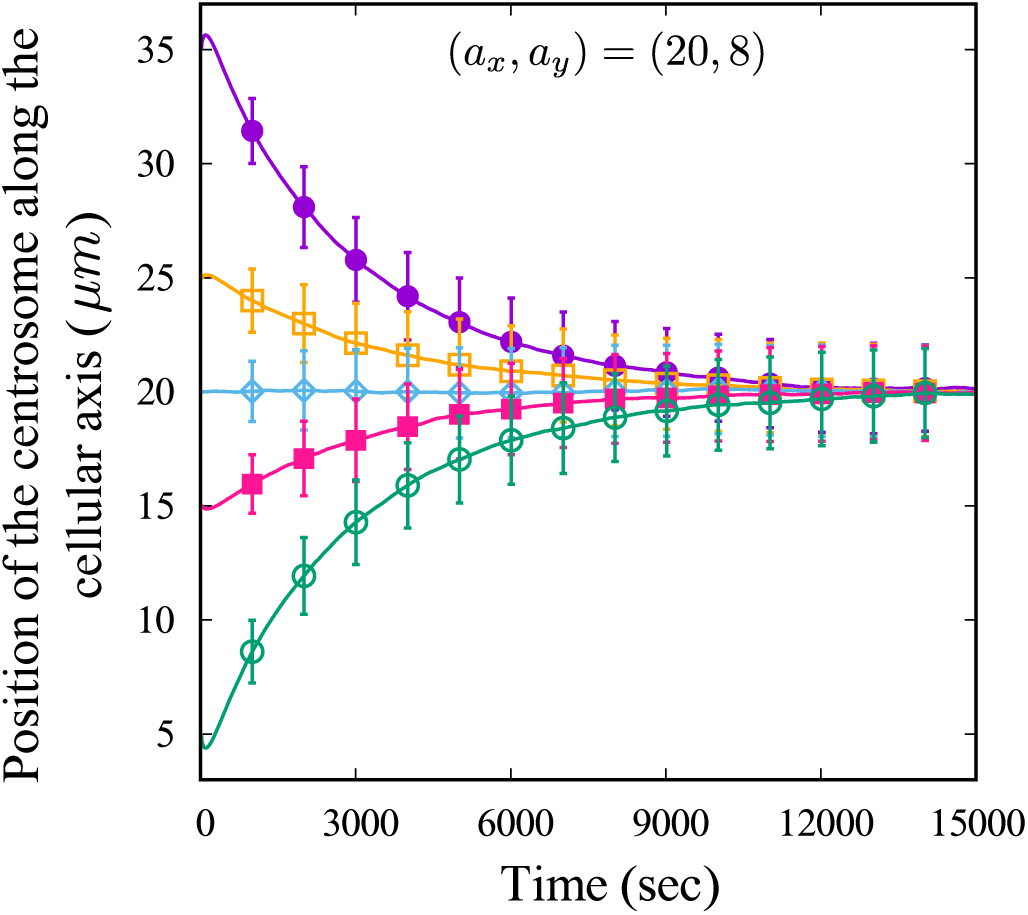
Centering of a single CS is achieved only with pulling forces; stoichiometric interaction between astral MTs and cortical dynein motors is enforced by fixing the number of cortical motors per astral MT interacting with the cortex.

## Acknowledgments

We thank Alex Mogilner for his careful readings of the manuscript and fruitful suggestions for its improvement. A.M. thanks the Indian Association for the Cultivation of Science (IACS), Kolkata, for support. A.S. was supported by a fellowship from the University Grants Commission (UGC), India. R.P. thanks IACS for support and Grant No. EMR/2017/001346 of SERB, DST, India for the computational facility.

